# An inositol pyrophosphate interaction screen provides insight into the regulation of plant casein kinase II

**DOI:** 10.1101/2025.11.27.691040

**Authors:** Kristina Sturm, Oded Pri-Tal, Felix Rico-Resendiz, Yash Verma, Annika Richter, Houming Chen, Larissa Broger, Ludwig A. Hothorn, Dorothea Fiedler, Vikram Govind Panse, Michael Hothorn

## Abstract

Inositol pyrophosphates (PP-InsPs) are key nutrient messengers in plants, but their protein receptors remain poorly defined. Using a systems-level affinity screen with biotinylated InsP₆, InsP₇, and InsP₈ in *Arabidopsis thaliana*, we identify multiple conserved PP-InsP-interacting complexes involved in mRNA metabolism, translation, and cell signaling, including the nuclear α-subunits of casein kinase II (CK2). The CK2 subunit AtCKA1 associates with the PP-InsP kinase AtVIH2, and its 1.9 Å crystal structure with InsP_6_ reveals two conserved PP-InsP binding sites located in the N-and C-terminal lobes. AtCKA1 binds InsP_6_, InsP_7_, and InsP_8_ with micromolar affinity. Mutation of both binding sites in the AtCKA^6xmut^ mutant abolishes PP-InsP binding in vitro. AtCKA^6xmut^ partially rescues the flowering phenotype of *ck2a1/2/3* mutants, and equivalent mutations inactivate the yeast orthologs ScCka1 and ScCka2. InsP_6_ competitively inhibits phosphorylation of canonical CK2 substrates by occupying a basic substrate-binding groove. Although incorporating β-subunits strongly enhances the phosphorylation of substrates by the AtCK2 holoenzyme, *ck2b1/2/3/4* mutants exhibit only mild growth defects in Arabidopsis. In *Marchantia*, loss of the single *ck2a* gene severely impairs growth, whereas deletion of the β subunit has no effect. Together, our findings suggest that InsP_6_/PP-InsPs modulate the activity of the isolated CK2 α-subunit by regulating access to its substrate-binding site.

## Introduction

Inositol pyrophosphates (PP-InsPs) are signaling molecules consisting of a fully phosphorylated myo-inositol ring and one or several pyrophosphate groups^1^. In plants, the inositol polyphosphate kinases, IPK1 and IPK2β catalyze the synthesis of inositol hexakisphosphate (InsP_6_)^2^. Inositol (1,3,4) trisphosphate 5/6-kinases (ITPKs) can subsequently convert InsP_6_ to InsP_7_^3–7^, and the diphosphoinositol pentakisphosphate kinases (PPIP5Ks) VIP/VIH then generate InsP_8_ from InsP_7_^8–12^. PP-InsP levels are tightly controlled in cells, in part by inositol pyrophosphatases^13–15^ that can be present at the C-terminus of PPIP5Ks^16,17^, or as stand-alone enzymes^18–22^.

The physiological and cellular functions for PP-InsPs in plants have been investigated in *atipk1-1 atipk2*β*-1* mutants, which lacked InsP_6_ and PP-InsPs, and which had elevated cellular levels of inorganic phosphate (Pi)^2,23^. *itpk* loss-of-function mutants and over-expression lines exhibit impaired Pi homeostasis in both Arabidopsis and *Marchantia polymorpha*^5,7,24,25^. In addition, auxin responses are altered in *atitpk1* plants^6^. Deletion of the PPIP5K AtVIH2 reduced cellular InsP_8_ pools and affected jasmonate-dependent defense responses^9^. Seedling lethal *atvih1 atvih2* double mutants had low InsP_8_ levels and hyperaccumulated Pi^11,26^. Deletion of the PPIP5K MpVIP1 in Marchantia resulted in reduced InsP_8_ pools, and in severe growth and developmental defects^20,27^. Mutations in inositol polyphosphate kinases altered salicylic acid (SA) accumulation and immune responses, highlighting a connection between PP-InsP metabolism and plant immunity^28–31^. Consistent with this, effector proteins possessing inositol polyphosphatase/pyrophosphatase activity have been identified^32,33^.

Thus far, few bona fide cellular receptors for PP-InsPs have been characterized in plants. The auxin receptor TRANSPORT INHIBITOR RESPONSE 1 (TIR1) requires InsP_6_ or a PP-InsP as essential co-factor^6,34^. Similarly, the jasmonate receptor CORONATINE-INSENSITIVE 1 (COI1) depends on a inositol polyphosphate co-factor^9,35–37^. In the context of Pi homeostasis and signaling^38^, SPX proteins function as cellular PP-InsP receptors^39^. Stand-alone plant SPX domains preferentially bind InsP_8_^26,40^, which promotes their interaction with PHOSPHATE STARVATION RESPONSE (PHR) transcription factors^41–45^. The resulting SPX – InsP_8_ – PHR complex prevents PHR1 from associating with its target promoters, thereby regulating Pi homeostasis and starvation responses^40,46,47^. InsP_8_ binding to the N-terminal SPX domain regulates the transport activity of the phosphate transporter AtPHO1^39,48–50^. Recently, a Marchantia DELLA protein has been characterized as both a MpVIP1-interacting protein and a direct PP-InsP receptor^27^.

Human casein kinase II (CK2) has been previously identified as an InsP_6_-binding protein^51–55^. CK2 is a ubiquitous eukaryotic serine/threonine protein kinase^56^ composed of two catalytic α-subunits and two regulatory β-subunits^57–59^. Substrates of CK2 frequently contain the consensus phosphorylation motif S/T-X-X-E/D^60^. A conserved basic surface patch adjacent to the CK2 active site engages with the acidic residues located downstream of the substrate phosphorylation site ^61^. Mutations in this basic patch alter CK2 substrate recognition^62^. The activity of the auto-catalytically active CK2 α-subunit can be modulated by the regulatory β-subunit, which contains acidic residues that are proposed to interact with a basic surface patch near the CK2α active site^62^. In addition, the β-subunit can directly interact with CK2 substrates^63^. CK2 exists either as an isolated CK2 α-subunit, as a α2β2 hetero-tetrameric holoenzyme^59^, or as a holoenzyme trimer^64,65^. Binding of InsP_6_ or InsP_7_ to the human CK2 α-subunit influences its interaction with specific protein substrates^53–55^. CK2 can also bind to and phosphorylate inositol hexakisphosphate kinases^66,67^. Furthermore, CK2 generates priming phosphorylation sites required for protein pyrophosphorylation, a post-translational modification mediated by PP-InsPs^67–70^.

Arabidopsis possesses three nuclear-localized CK2 α-subunit isoforms (AtCKA1, AtCKA2, AtCKA3), while a fourth isoform, AtCKA4, contains a transit peptide that directs its localization to the chloroplast^71,72^. Additionally, four CK2 β-subunits (AtCKBs) are localized to the cytosol and/or nucleus^71^. Constitutive expression of an inactive CK2 α-subunit from tobacco in Arabidopsis induced a strong dominant-negative effect on plant growth, development, and cell cycle progression ^73^, and further disrupted DNA repair, SA biosynthesis, and auxin transport^74–76^. By contrast, *cka1/2/3* mutants displayed comparatively mild phenotypes, including reduced hypocotyl elongation, smaller cotyledons, decreased lateral root formation, delayed flowering, and altered circadian rhythms^77,78^. Functional knockdown of CK2A4 in the *cka1/2/3* mutant background suggested that all four α-subunits act redundantly in regulating lateral root development and flowering time^72^. Chemical inhibition of CK2 α-subunits by the small molecule CK2 inhibitor TTP-22 reduced primary root growth and lead to increased cellular Pi levels^79^. Single *cka4* T-DNA mutants exhibited reduced primary root length, enhanced lateral root density, delayed cotyledon expansion, and flowering^80^. Overexpression of specific CK2 β-subunits induced early flowering, resulted in shorter hypocotyls under far-red light conditions, and altered circadian rhythms^81–83^.

Similar to other eukaryotes^60^, plant CK2 phosphorylates a wide range of substrates, thereby regulating different signaling pathways. Among many transcription factors regulated by Arabidopsis CK2 is the Myb-domain protein CIRCADIAN CLOCK ASSOCIATED 1 (AtCCA1), whose DNA-binding activity is stimulated by CK2-dependent phosphorylation^81,84,85^. Another key target is SUPPRESSOR OF GAMMA RADIATION1 (AtSOG1), a functional homolog of the mammalian tumor suppressor p53 and a central regulator of plant DNA damage responses^86–88^. AtCK2 phosphorylates AtSOG1, and together these proteins coordinate seedling root growth under Pi starvation^79^. In rice, OsCK2 also modulates Pi homeostasis by phosphorylating the phosphate transporter OsPHT1, leading to its retention in the endoplasmic reticulum^89^.

In this study, we report a system-wide screen for InsP_6_/PP-InsP interacting proteins in Arabidopsis, which reveals several novel, putative InsP_6_/PP-InsP receptors involved in mRNA metabolism, translation and cell signaling. Among the candidates are all three nuclear-localized AtCK2A α-subunits.

## Results

### A small-molecule interaction screen yields novel candidate PP-InsP receptors

To identify candidate PP-InsP–interacting proteins, we incubated protein extracts from Arabidopsis seedlings grown under phosphate (Pi) starvation with streptavidin-coupled sepharose beads functionalized with biotinylated InsP_6_ or non-hydrolyzable analogs of InsP_7_ and InsP_8_^90–92^. The InsP_6_/PP-InsP analogs were immobilized via their 1/3′ position, leaving the axial 2′ phosphate group accessible for interaction with target proteins in the presence of EDTA (Supplementary Fig. 1a). Bound proteins were subsequently eluted from the beads by competition with excess soluble InsP_6_, 5PCP-InsP_5_ (hereafter referred to as InsP_7_) and 1,5(PCP)_2_-InsP_4_ (hereafter referred to as InsP_8_) (Supplementary Fig. 1b)^90^. Label-free quantification revealed ∼600 potential interactors (Supplementary Table 1). Specifically, we identified 406 proteins in the InsP_6_ dataset, 526 in the InsP_7_ dataset, and 456 in the InsP_8_ dataset. Considerable overlap was observed across conditions, with 269 proteins (45% of the total hits) shared among all three datasets, consistent with patterns recently reported for the human PP-InsP interactome^92^ (Fig. 1a; Supplementary Table 1). Furthermore, the Arabidopsis InsP_7_ dataset displayed substantial overlap with our previously characterized InsP_7_ interactomes from yeast and human cells (Fig. 1b) ^91,93^. We combined the hits from the Arabidopsis InsP_6_, InsP_7_ and InsP_8_ samples (Supplementary Fig. 1c-e) for the subsequent analysis, unless indicated otherwise.

**Fig. 1:**
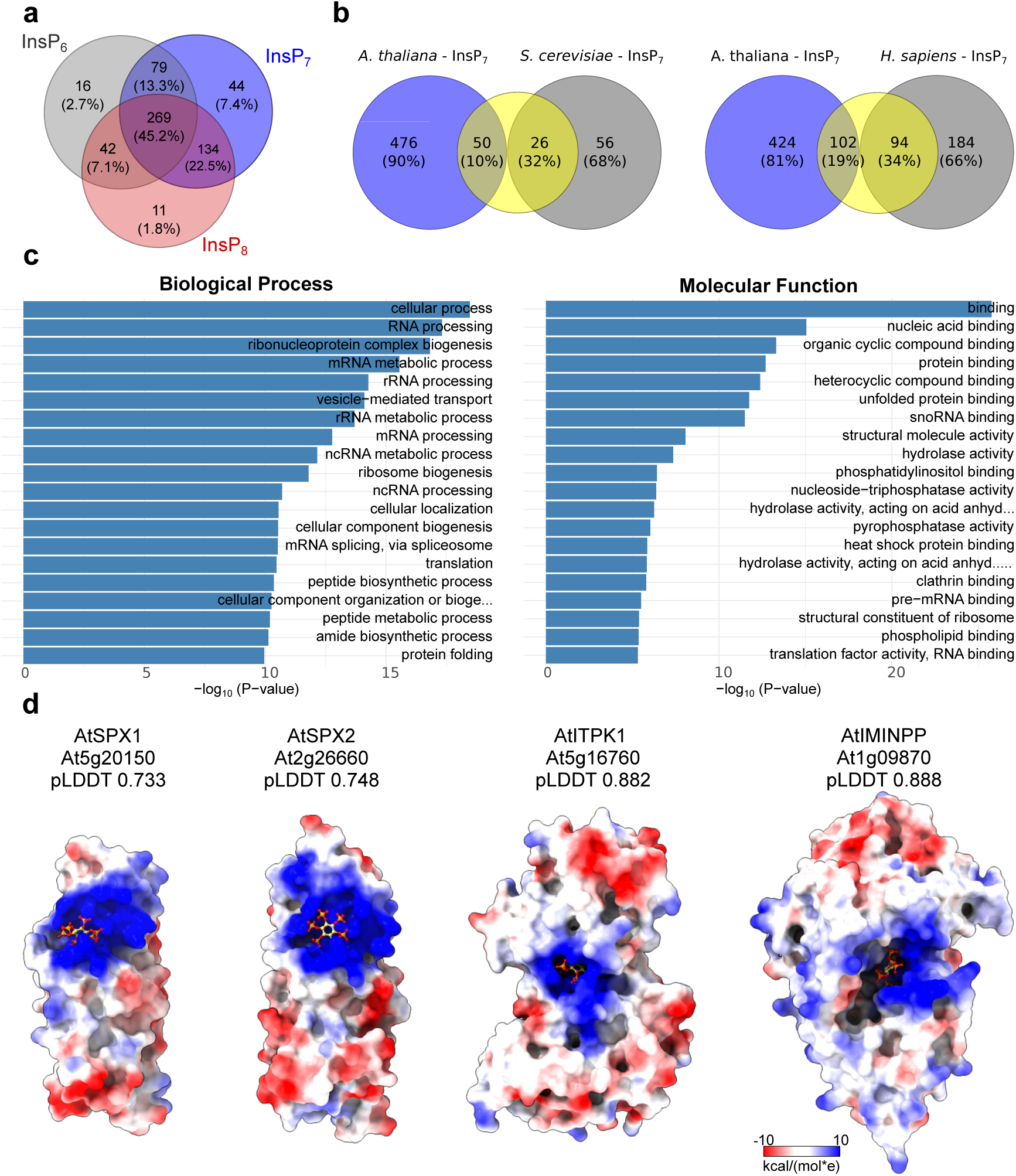
A InsP_6_ / PP-InsP affinity screen yields a large number of potential interactors in Arabidopsis. **a** Venn diagram comparing the number of proteins enriched in the 1/3-biotin-InsP_6_ (InsP_6_, in gray), 1/3-biotin-5PCP-InsP_5_ (InsP_7_, in blue) or 3-biotin-1,5-(PCP)_2_-InsP_4_ (InsP_8_, in red) samples. **b** Venn diagrams comparing the number of proteins enriched in the InsP_7_ sample (in blue) compared to sequence-related proteins previously identified in the yeast (left) and human InsP_7_ (right) interactomes (in gray). Sequence homologs were identified in reciprocal blastp searches with an e-value cutoff of 1e−5 (yellow circle). **c** Gene ontology (GO) analysis for the 594 proteins identified across the InsP_6_, InsP_7_ and InsP_8_ samples. **d** Surface views of the known PP-InsP binding proteins AtSPX1 (TAIR-ID: At5g20150, https://www.arabidopsis.org; AF2-ID: AF-A0A654G2V5-F1-v4, https://alphafold.ebi.ac.uk/), AtSPX2 (At2g26660, AF-A0A5S9X1N0-F1-v4), AtITPK1 (At5g16760, AF-A0A178U6K3-F1-v4) and AtMINPP (At1g09870, AF-Q941B2-F1-v4). Shown are electrostatic surface potentials in kcal/(mol*e) at 298 K from –10 (red) over 0 (white) to 10 (blue), mapped to AlphaFold2 models; docked InsP_6_ molecules (in bonds representation, AtSPX1 Boltz-2 pIC_50_ 570 nM, AtSPX2 pIC_50_ 1.1 μM, AtITPK1 pIC50 1.7 μM, AtMINPP pIC50 1.9 μM).

Gene Ontology (GO) analysis of the combined dataset revealed strong enrichment for processes related to RNA metabolism and ribosome biology (Fig. 1c). We recovered several known plant PP-InsP-binding proteins, including AtSPX1, AtSPX2^26,39,40^, the inositol polyphosphate kinases AtIPK2β^2^ and AtITPK1^3,94^ and the MULTIPLE INOSITOL POLYPHOSPHATE PHOSPHATASE (AtMINPP) (Supplementary Table 1). These proteins are all characterized by a large basic surface patch that facilitates InsP_6_/PP-InsP coordination (Fig. 1d). Other known Arabidopsis InsP_6_/PP-InsP-binding proteins including AtTIR1^34^, AtCOI1^9,35,37^ and AtPHO1^39,49^ were absent from our interactome (Supplementary Table 1).

We next aimed to identify new candidate InsP_6_/PP-InsP binding proteins. To date, approximately 300 protein structures with bound InsP_6_ have been deposited in the Protein Data Bank (PDB, http://rcsb.org, chemical ID: IHP). We identified clear sequence homologs for about half of them within our InsP_6_/PP-InsP interactome (Fig. 2a). Functional annotation of this subset of our plant interactome revealed significant enrichment in processes related to mRNA processing and splicing (Fig. 2b). Consistent with this, InsP_6_ was previously identified as being bound to the subunit Prp8 in cryo-electron microscopy (cryo-EM) structures of fungal and animal spliceosomes^95,96^. Notably, PRP8 was recovered in our InsP_6_ sample (Supplementary Fig. 1c), as well as in the previously reported the yeast and human InsP_7_ interactomes (Fig. 1b)^91,93^. Molecular docking confirmed that the conserved inositol polyphosphate binding pocket of AtPRP8A can accommodate an InsP_6_ molecule (Fig. 2c). From the 61 protein subunits in the human pre-catalytic spliceosome precursor^97^, 47 were present in our interactome and could be mapped onto the complex structure (Fig. 2d; Supplementary Table 1). Together, these findings suggest that fungal, plant and human spliceosomal complexes associate with InsP_6_ and/or PP-InsPs in vivo, likely through the conserved PRP8 subunit. Supporting this notion, suppressor mutations map to the inositol polyphosphate binding site of yeast Prp8^98^.

**Fig. 2:**
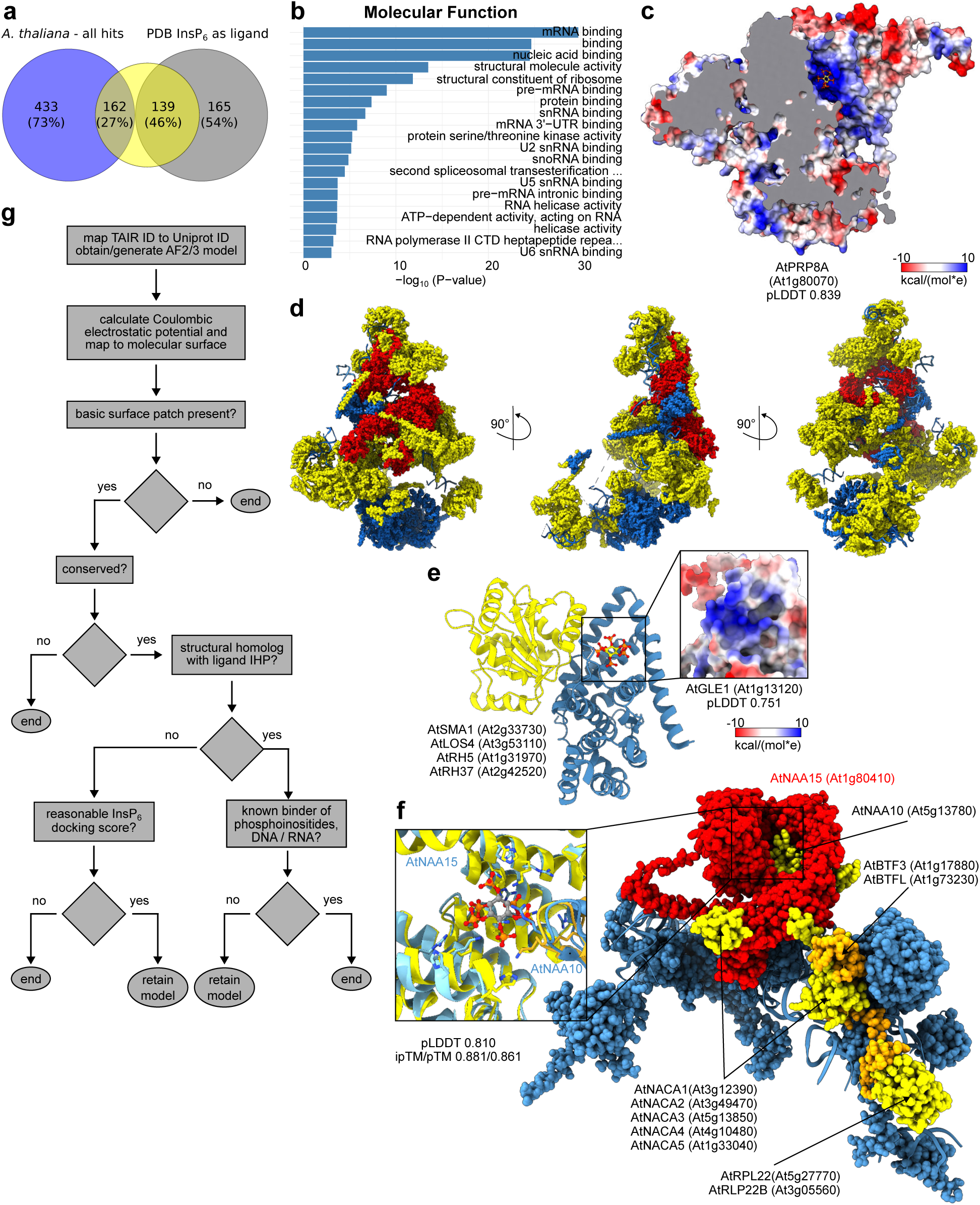
Structure-guided identification of putative InsP_6_/PP-InsP binding protein complexes. **a** Venn diagram comparing proteins enriched in the InsP_6_/PP-InsP pull down (in blue) with all crystal and single-particle cryo-electron microscopy structures containing InsP_6_ (chemical ID: IHP, September 2025, in gray) in the Protein Data Bank (PDB, https://rcsb.org). Sequence homologs were identified in reciprocal blastp searches with an e-value cutoff of 1e−5 (yellow circle). **b** Gene Ontology (GO) analysis of 162 interactome proteins exhibiting significant sequence similarity to InsP₆-bound proteins (or protein complexes) reported in the PDB. **c** Electrostatic surface potential mapped onto the AlphaFold2-predicted structure of *Arabidopsis thaliana* PRP8A (At1g80070, AF-Q9SSD2-F1-v4, pLDDT value indicated), with a docked InsP₆ (Boltz-2 pIC_50_ 2.4 μM) molecule shown in bond representation. **d** Structure of the human pre-catalytic spliceosome precursor (PDB-ID: pdb_00006ah0, in blue), with the Prp8 subunit highlighted in red and 47 additional subunits from the interactome shown in yellow. **e** Ribbon diagram of the Dbp5 (yellow) – Gle1 (blue) – InsP₆ (bonds representation) complex (PDB-ID: pdb_00003peu). Four Dbp5-like RNA helicases identified in the screen are shown alongside. Inset: molecular surface view of the predicted InsP₆/PP-InsP binding pocket in AtGle1 (At1g13120, AF-Q0WPZ7-F1-v4). **f** Structure of the human NatA –NAC – MAP1 80S ribosome complex (PDB-ID: pdb_00009fq0) with the InsP_6_-coordinating Naa15 subunit shown in red and four additional complex subunits present in the interactome highlighted in yellow. Inset: structural superposition of HsNaa15 (in yellow) and AtNAA15 (in blue, AF-Q8VZM1-F1-v4, r.m.s.d. is ∼0.8 Å comparing 461 corresponding C_α_ atoms), highlighting the conservation of the InsP_6_ binding site. Short hairpins from HsNaa10 (yellow) and AtNAA10 (blue) that complete the InsP₆-binding pocket are shown alongside. **g** Flowchart summarizing the structural bioinformatic pipeline used to analyze ∼600 candidates from the interactome.

A second family of mRNA processing factors recovered from our InsP_6_/PP-InsP interactome are RNA helicases involved in mRNA protein complex remodeling. It has been previously shown that the ATPase activity of the yeast RNA helicase Dbp5 is stimulated by InsP_6_ and that InsP_6_ acts as a molecular glue promoting the interaction of Dbp5 with the mRNA export factor Gle1^99,100^. We recovered four Dbp5-like RNA helicases in our interactome and found that the InsP_6_ basic binding surface is also present in AtGLE1 (Fig. 2e). This suggest that InsP_6_ or a PP-InsP could be involved in the regulation of AtGLE1 mediated mRNA export in Arabidopsis.

Among the enriched GO terms in our analysis is ‘structural constituent of ribosome’ (Figs. 1c; 2b). In our screen, we identified the catalytic and auxiliary subunits AtNAA10 and AtNAA15 of the N-terminal acetyltransferase A (AtNatA) complex (Supplementary Fig. 1c-e; Supplementary Table 1), an enzyme that transfers an acetyl group from acetyl-CoA to the N-terminus of nascent polypeptides as they emerge from the ribosome^101,102^. Yeast and human NatA complexes bind InsP_6_ with nanomolar affinity at the Naa10 – Naa15 interface^103,104^, where InsP_6_ is coordinated by a set of basic residues also present in AtNAA15 and AtNAA10 (Fig. 2f). A recent cryo-EM study revealed the structure of the human NatA complex bound to the 80S ribosome^105^. Our InsP_6_/PP-InsP interactome includes ribosomal proteins and ribosome-associated factors, many of which could be confidently mapped to the NatA – 80S complex. These include all five isoforms of the nascent polypeptide-associated complex (NAC; AtNACA1-5), both isoforms of basic transcription factor 3 (AtBTF3), and two isoforms of ribosomal protein L22 (AtRLP22) (Fig. 2f). Together, these findings suggest that the AtNatA complex interacts with InsP_6_/PP-InsPs in planta while bound to the 80S ribosome, thereby rationalizing the presence of ∼50 ribosomal proteins and ribosome-associated factors in our interactome (Supplementary Table 1).

We next aimed to uncover additional, less abundant InsP_6_/PP-InsP interacting proteins. Known InsP_6_/PP-InsP binding sites are characterized by clusters of conserved, surface-exposed basic residues (Figs. 1d; 2c-f). To systematically search for such features, we obtained AlphaFold2 ^106^ structural models for the ∼600 candidate proteins identified in our interactome and analyzed them for the presence of conserved basic surface patches within globular domains (Fig. 2g). This approach revealed additional putative inositol polyphosphate binding proteins with structural homologs bound to InsP_6_ in the PDB. Among the candidate proteins is the cohesin-associated subunit Pds5, which plays a critical role in the loading and release of cohesin on chromatin^107^. InsP_6_ has been characterized as a structural co-factor of Pds5, stabilizing its conformation and facilitating its interaction with cohesin^108^ (Fig. 3a). Within our interactome, we detected the Arabidopsis homolog AtPDS5A, which shares 25% sequence identity with human Pds5 (Supplementary Fig. 1d,e; Supplementary Table 1). Structural analyses of human Pds5 revealed that InsP_6_ binds to a large basic surface patch located in the C-terminal lobe of the HEAT repeat domain ^108^, a feature that is highly conserved in AtPDS5A (Fig. 3a).

**Fig. 3:**
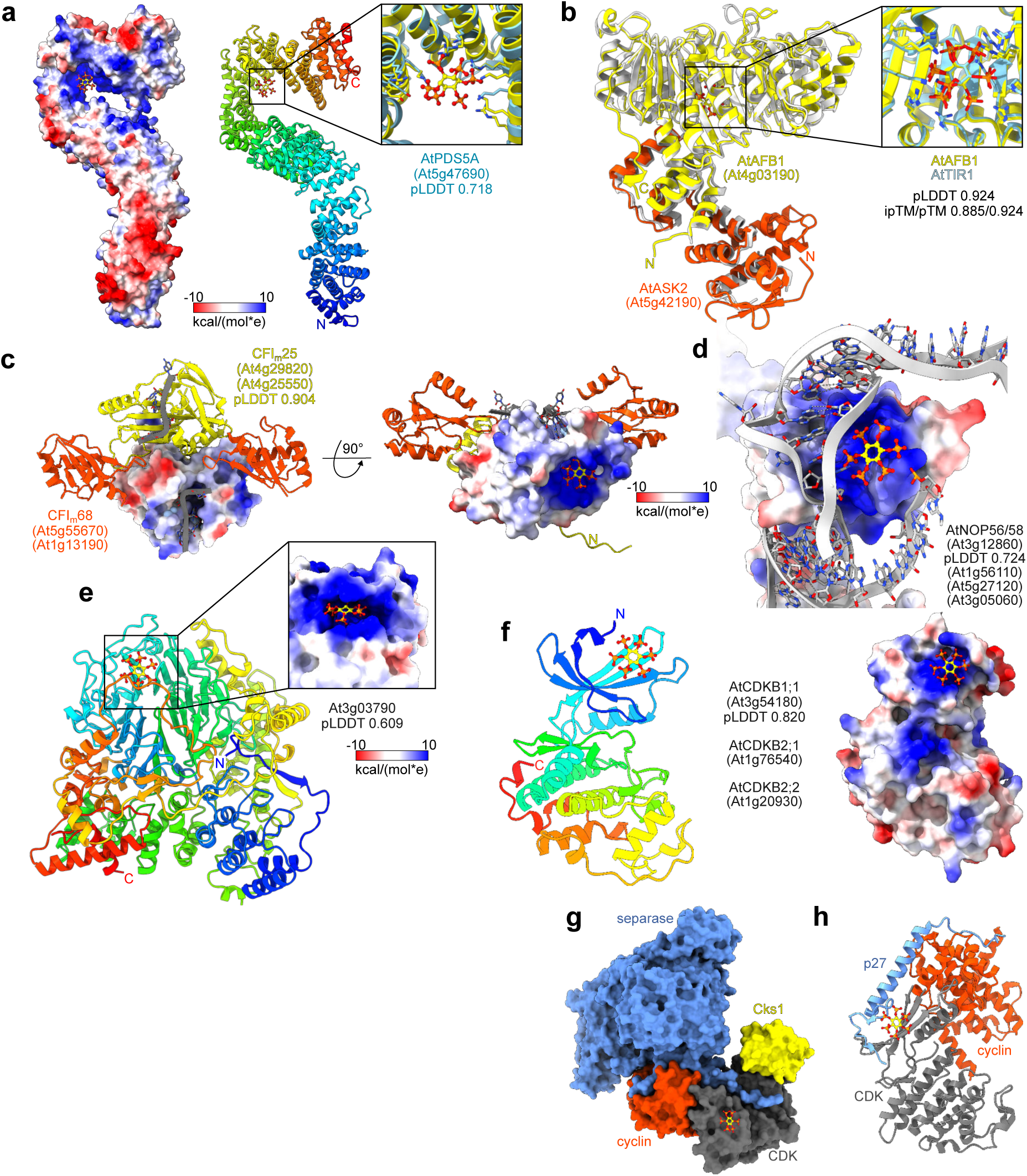
Protein complexes involved in hormone signaling, mRNA/rRNA processing and cell cycle control bind InsP_6_/PP-InsPs in planta. **a** Electrostatic surface potential mapped onto the crystal structure of human Pds5 (PDB-ID: pdb_00005hdt) bound to InsP_6_ (in bonds representation), a ribbon diagram colored from blue (N-term) to red (C-term) is shown alongside. Inset: structural superposition of the C-terminal HsPds5 HEAT repeats (in yellow) with the corresponding region in AtPDS5A (in blue, AlphaFold3 model, root mean square deviation is ∼1.2 Å comparing 245 corresponding C_α_ atoms) reveals the InsP_6_ binding surface to be well conserved. **b** Structural superposition of the Alphafold3 modeled AtAFB1 (yellow) – AtASK2 (orange) auxin receptor complex with the crystal structure of AtTIR1 – AtASK1 (PDB-ID pdb_00002p1m, in gray, r.m.s.d. is ∼0.8 Å comparing 566 corresponding C_α_ atoms). Inset: View of the putative InsP_6_ binding surface in AtAFB1 (in yellow, in bonds representation, Boltz-2 pIC_50_ is 5.0 μM) compared to the AtTIR1 – InsP_6_ complex (in blue). **c** Structure of the RNA-bound CFI_m_ complex (PDB-ID: pdb_00002q2t) with CFI_m_25 and CFI_m_68 shown in yellow and orange, respectively, and including an electrostatic surface potential mapped onto the AlphaFold3 model of AtCFI_m_25 (At4g25550, r.m.s.d. is ∼0.8 Å comparing 185 corresponding C_α_ atoms). The docked InsP_6_ molecule is shown in bonds representation (Boltz-2 pIC_50_ 5.3 μM). The RRM domains of the putative CFI_m_68 subunits At5g55670 and At1g13190 superimpose with r.m.s.d.’s of ∼1 Å comparing 75 corresponding C_α_ atoms. **d** Electrostatic surface potential mapped onto the AlphaFold3 model of AtNOP56 (At3g12860) and superimposed with the human NOP56 subunit within the human small ribosomal subunit processome (PDB-ID: pdb_00007mq8, r.m.s.d is ∼1.1 Å comparing 205 corresponding C_α_ atoms, Boltz-2 pIC_50_ 4.1 μM). A fragment of the U3 rRNA is shown alongside (in gray). **e** Ribbon diagram of the Alphafold3 model of At3g03790 with a docked InsP_6_ molecule (in bonds representation, Boltz-2 pIC_50_ 300 nM) Inset: Electrostatic surface potential of the putative InsP_6_ binding surface. **f** (left panel) Ribbon diagram colored from blue (N-term) to red (C-term) of AtCDKB2;1 (At1g76540, AF-A0A178WJF2-F1-v4) with a docked InsP_6_ molecule (in bonds representation, Boltz-2 pIC_50_ 6.3 μM). (right panel) Electrostatic surface potential highlighting the basic surface patch in the N-lobe of AtCDKB2;1. **g** Surface representation of the human separase (in blue) – Cdk1 (in gray – cyclin B1 (in orange) – Cks1 (in yellow) complex (PDB-ID pdb_00007nj0) and including the position of the putative InsP_6_ binding site (in bonds representation). **h** Ribbon diagram of a Cdk4 (in gray) – cyclin D1 (in orange) – p27 complex (PDB-ID: pdb_00006p8e, in blue) with the relative position of the InsP_6_ molecule indicated (in bonds representation).

AUXIN-SIGNALING F-BOX 1 (AFB1) functions as a rapid-response auxin receptor^109,110^. In our screen, we identified AtAFB1 together with AtSKP1B/ASK2 and constructed a structural model of the heterodimeric complex bound to InsP_6_ using AlphaFold3^111^ and Boltz-2^112^ (Fig. 3b). The predicted complex aligns closely with the experimentally determined AtTIR1 – AtSKP1A – InsP_6_ structure^34^. The docked InsP_6_ molecule occupies a position nearly identical to that observed in the crystal structure, indicating that the docking approach can reliably identify candidate InsP_6_/PP-InsP binding sites (Fig. 3b).

Building on this rationale, we broadened our analysis to encompass interactome-derived candidates that lacked an experimentally resolved InsP_6_-bound homolog in the PDB, but exhibited a conserved basic surface patch compatible with inositol polyphosphate coordination. Through this approach, we uncovered additional factors associated with mRNA processing, specifically subunits of the cleavage factor Im (CFI_m_) complex, which plays a central role in polyadenylation site selection. Human CFI_m_25 is a metabolite– and RNA-binding protein with a Nudix-fold enzyme architecture but devoid of catalytic activity^113,114^. Both Arabidopsis CFI_m_25 isoforms identified in our interactome (Supplementary Table 1) have a conserved basic surface patch distinct from the known RNA recognition interface (Fig. 3c). Molecular docking analyses indicated that this patch is capable of accommodating InsP_6_ or a PP-InsP (Fig. 3c). Given that CFI_m_25 forms a heterotetrameric complex with the CFI_m_68 subunit through their RRM RNA-binding domains^114^, and that we identified two putative Arabidopsis CFI_m_68 homologs in our screen (Supplementary Table 1), it is likely that InsP_6_ or a PP-InsP may engage with an intact CFI_m_ complex in planta (Fig. 3c).

Small nucleolar RNA (snoRNA)-associated proteins that function in ribosomal RNA processing^115,116^, were found to be enriched in our interactome (Fig. 1c). Within this protein set, we identified all four Arabidopsis NOP56/58 isoforms (Supplementary Fig. 1c-e; Supplementary Table 1), each of which harbors a conserved basic surface patch able to accommodate an InsP_6_ or PP-InsP molecule (Fig. 3d). In human NOP56, the corresponding surface region is not involved in U3 snoRNA binding within the maturing small subunit processome during ribosome biogenesis (Fig. 3d)^117^. The identification of putative InsP_6_/PP-InsP binding sites in snoRNA-associated proteins may partly explain the enrichment of nucleolar and ribosomal proteins in our dataset (Fig. 1c) and in the yeast and human interactomes^91–93^.

At3g03790 identified in our InsP_6_/PP-InsP interaction screen, encodes a single-copy gene of unknown function in Arabidopsis. The predicted protein harbors N-terminal ankyrin repeat motifs and a WD40 domain that exhibits structural similarity to the UV-B photoreceptor UVR8^118^ (Fig. 3e). Within the WD40 domain, a prominent central basic surface patch is present, which is distinct from the interaction surfaces previously described for WD40-containing RNA-binding proteins^119^. Molecular docking experiments with Boltz-2 suggest that this basic surface patch in At3g03790 may be able to accommodate an InsP_6_/PP-InsP ligand (Fig. 3e).

Finally, several protein kinase families were represented in our InsP_6_/PP-InsP interactome. These include AtCDKB1;1, AtCDKB2;1, and AtCDKB2;2, members of the B-type cyclin-dependent kinases (CDKBs) that function in cell cycle regulation^120^. All isoforms contain a conserved basic surface patch within the N-lobe of the kinase domain, to which an InsP_6_ molecule could be docked (Fig. 3f). This basic binding surface is also present in human Cdk1 and Cdk2, which together with cyclin B1 are part of the human InsP_7_ interactome^91^. The putative InsP_6_/PP-InsP binding site remains accessible in the Cdk1 – cyclin B1 – Cks1 – separase complex (Fig. 3g)^121^, yet it is positioned near the binding site of the p27 regulatory protein in the human Cdk4 – cyclin D1 – p27 complex (Fig. 3h)^122^. This suggests that InsP_6_/PP-InsPs may target plant and animal cyclin-dependent kinases to modulate their interaction with p27-like inhibitors, which are also present in Arabidopsis^120^.

Overall, our comparative sequence and structural analyses of plant, fungal and human InsP_6_/PP-InsP interactomes indicate the existence of previously unrecognized, evolutionarily conserved PP-InsP binding proteins that have molecular roles in RNA biology, protein synthesis, and cell signaling.

### Nuclear-localized Arabidopsis casein kinase II α-subunits bind InsP_6_/PP-InsP

Previous studies have demonstrated that PP-InsP binding proteins physically interact with PP-InsP-biosynthetic enzymes^27,66,67^. In order to identify bona fide PP-InsP receptors among the numerous InsP_6_/PP-InsP binding proteins in Arabidopsis (see above), we screened for interactors of AtVIH2, a PPIP5K that catalyzes the conversion of InsP_7_ to InsP_8_, using immunoprecipitation followed by mass spectrometry (IP/MS). For this, we employed the previously reported functional pAtVIH2::AtVIH2-mCit fusion protein expressed in the *vih1-2 vih2-4* mutant background^11^. We identified 472 putative VIH2-interacting proteins (Fig. 4a; Supplementary Table 2). Of these, 82 were also present in our InsP_6_/PP-InsP interactome (Fig. 4a). Among the candidates were known or putative InsP_6_/PP-InsP-associated proteins, including AtPRP8A (Fig. 2c), AtLOS4 (Fig. 2e), AtNAA15 (Fig. 2f), AtPDS5A (Fig. 3a), and the casein kinase II α-subunit isoform 1 (AtCKA1) (Fig. 4b). We confirmed that the nuclear-localized AtCKA1, but not the plastid-localized AtCKA4 (which shares 85% sequence identity with AtCKA1), interacts with AtVIH2, by performing split-luciferase assays in *Nicotiana benthamiana* (Fig. 4c). Notably, all three nuclear-localized CK2 α-subunits AtCKA1, AtCKA2 and AtCKA3 are present in our InsP_6_/PP-InsP interactome (Supplementary Fig. 1c-e; Supplementary Table 1).

**Fig. 4:**
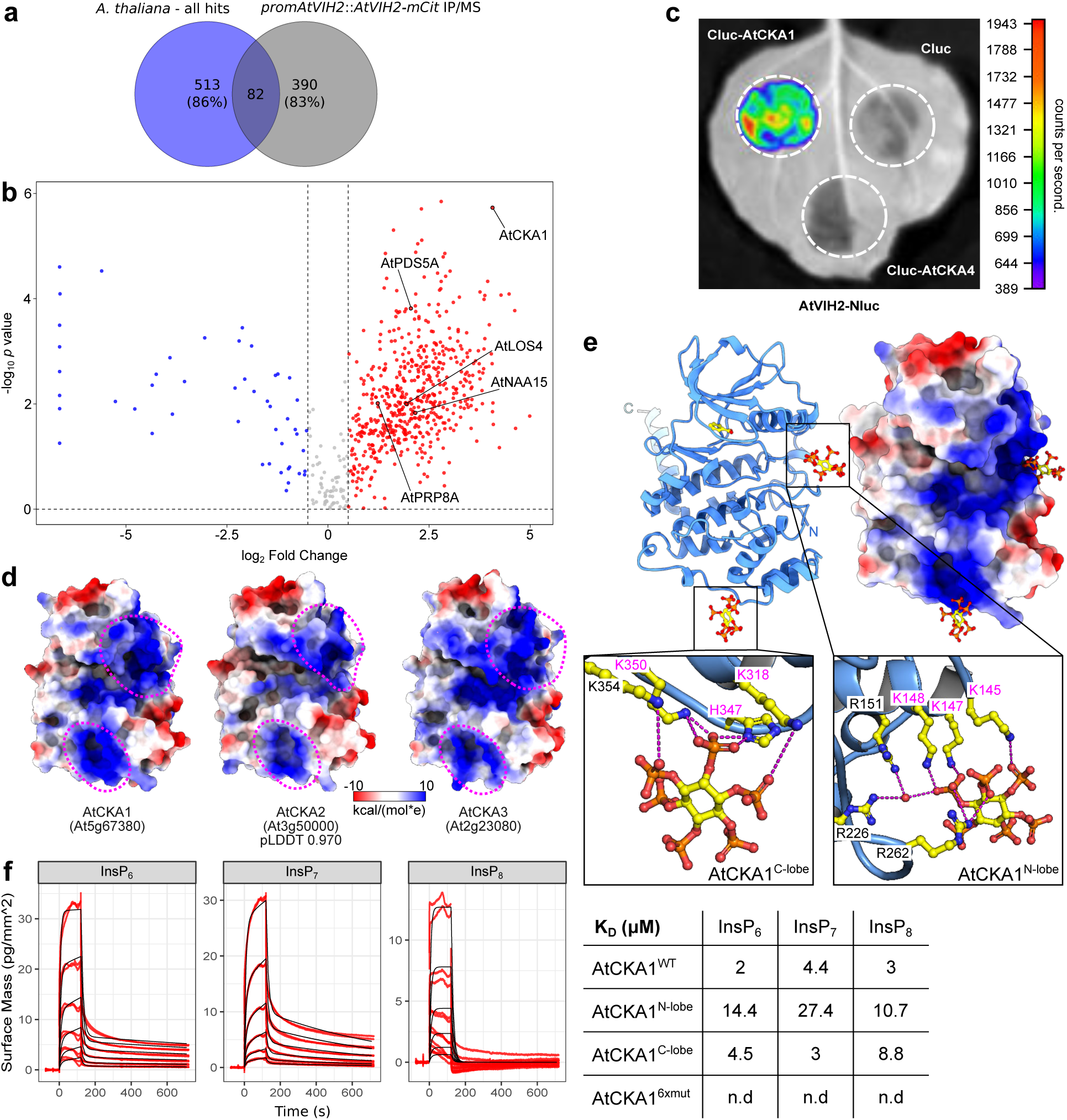
Nuclear-localized CK2 α-subunits sense InsP_6_/PP-InsPs via two conserved binding surfaces. **a** Venn diagram comparing proteins enriched in the InsP₆/PP-InsP pull down (blue) with putative AtVIH2 interacting proteins identified by IP/MS (gray). **b** Volcano plots showing significantly enriched proteins (FDR < 0.05) in the AtVIH2-mCit immunoprecipitate compared to the YFP control. Selected proteins discussed in the text are indicated. **c** Split-luciferase assay in *N. benthamiana* leaves between AtVIH2 and AtCKA1. Cluc-AtCKA4 and a Cluc empty vector were used as controls. The color bar shows the arbitrary signal intensity with pseudo color applied. **d** Structural comparison of the three nuclear CK2A isoforms. Shown are electrostatic surface potentials in kcal/(mol*e) at 298 K from –10 (red) over 0 (white) to 10 (blue), mapped to the crystal structure of AtCKA1 (PDB-ID: pdb_00006xx6, left)^123^, to the Alphafold2 model of AtCKA2 (AF-Q08466-F1-model_v4, center), and to the 1.25 Å resolution structure of AtCKA3 (right). The basic surfaces patches adjacent to the nucleotide-binding cleft and the C-terminal lobe are highlighted with dotted lines (in magenta). **e** Crystal structure of the AtCKA1 – InsP_6_ complex. Shown are a ribbon diagram (in blue, left) and an electrostatic surface potential (right). The InsP _6_ ligand is highlighted in bonds representation, a benzoic acid molecule occupying the nucleotide binding cleft is shown alongside (in yellow). The insets provide details of the InsP_6_/PP-InsP binding surfaces. Conserved interacting residues are highlighted in bonds representation, hydrogen bond interactions, in part mediated by a water molecule (red sphere), are indicated by dotted lines (in magenta). **f** GCI analysis of ligand-binding kinetics between AtCKA1 and 1/3-biotin-InsP_6_ (InsP_6_), 1/3-biotin-5PCP-InsP_5_ (InsP_7_) and 3-biotin1,5(PCP)_2_-InsP_4_ (InsP_8_) coupled to the GCI chip. Shown are sensorgrams (n=2, red) with global fits (black). The table provides the equilibrium dissociation constants (K_D_) for the wild-type (WT) kinase, the K145A/K147A/K148A mutant (N-lobe), the K318A/H347A/K350A mutant (C-lobe) and the K145A/K147A/K148A/K318A/H347A/K350A mutant (6xmut).

We compared the previously determined structure of AtCKA1^79,123^ with the Alphafold2 model of AtCKA2 and a crystal structure of apo AtCKA3 refined to 1.25 Å resolution (Fig. 4d; Supplementary Table 3). All three isoforms harbor two conserved basic surface patches, located in close proximity to the nucleotide-binding cleft of the kinase and at the C-terminal lobe of the enzyme, respectively (Fig. 4d). A 1.9 Å structure of AtCK2A1 crystallized in the presence of InsP _6_ revealed an InsP_6_ molecule bound at the interface of two adjacent kinase molecules, and occupying two potential binding sites (Fig. 4e; Supplementary Fig. 2a). One binding site is formed by the basic patch adjacent to the nucleotide-binding cleft, while the second site maps to the C-terminal lobe of the enzyme (Fig. 4e). Both binding sites are also found in contact with an InsP_6_ molecule in the previously reported structure of human CK2 (Supplementary Fig. 2b)^54^. Conserved lysine and histidine residues form hydrogen bond interactions with the phosphate groups of the ligand, in part mediated by water molecules (Fig. 4e; Supplementary Fig. 3).

We next quantified the interaction of AtCKA1 with InsP_6_/PP-InsPs using grating-coupled interferometry (GCI). Wild-type AtCKA1 bound InsP_6_, InsP_7_ and InsP_8_ with low micromolar affinity (Fig. 4f). Mutation of three conserved lysine residues in the binding surface adjacent to the nucleotide-binding cleft (AtCKA1^N-lobe^, K145A/K147A/K148A) reduced InsP_6_/PP-InsP binding by approximately four to seven-fold (Fig. 4f; Supplementary Fig. 4). Mutation of the binding surface at the C-terminal lobe of the kinase (AtCKA1^C-lobe^, K318A/H347A/K350A) had little effect (Fig. 4f; Supplementary Fig. 4). However, simultaneous mutation of both binding surfaces (AtCKA1^6xmut^) abolished binding to InsP_6_, InsP_7_ and InsP_8_ (Fig. 4f; Supplementary Fig. 4).

Taken together, our proteomic, structural, and quantitative biochemical experiments identify the three nuclear-localized α-subunits of casein kinase II as bona fide InsP_6_/PP-InsP binding proteins in Arabidopsis.

### InsP_6_ targets the CK2 substrate binding site

We hypothesized that the InsP_6_/PP-InsP binding sites, one of which is located in close proximity to the nucleotide-binding cleft of AtCKA1, might affect the enzyme’s interaction with its protein substrates. We purified fragments of two known AtCKA1 substrates, AtCCA1^81,84,85^ and AtSOG1^79^, and performed in vitro kinase assays. Bovine milk α-casein was included as a generic CK2 substrate. Wild-type AtCKA1 robustly phosphorylated AtSOG1, AtCCA1, and casein (Fig. 5a). Phosphorylation of all three substrates was markedly reduced in the presence of 200 μM InsP_6_; the previously reported ATCKA1 kinase inhibitor TTP-22^79^ had a similar effect (Fig. 5a). The InsP_6_/PP-InsP binding site mutant AtCKA1^6xmut^ exhibited reduced trans-phosphorylation activity toward AtSOG1 and casein (Fig. 5b). Notably, the addition of InsP_6_ did not further reduce phosphorylation of protein substrates by the AtCKA1^6xmut^, which lacks detectable InsP_6_ binding (Figs. 5b; 4f). However, the mutant kinase remained sensitive towards to the active-site inhibitor TTP-22 (Fig. 5b). Together, these experiments suggest that InsP_6_ can bind to the InsP_6_/PP-InsP interaction surfaces identified in the AtCKA1 – InsP_6_ complex structure and reduces AtCKA1’s ability to trans-phosphorylate protein substrates.

**Fig. 5:**
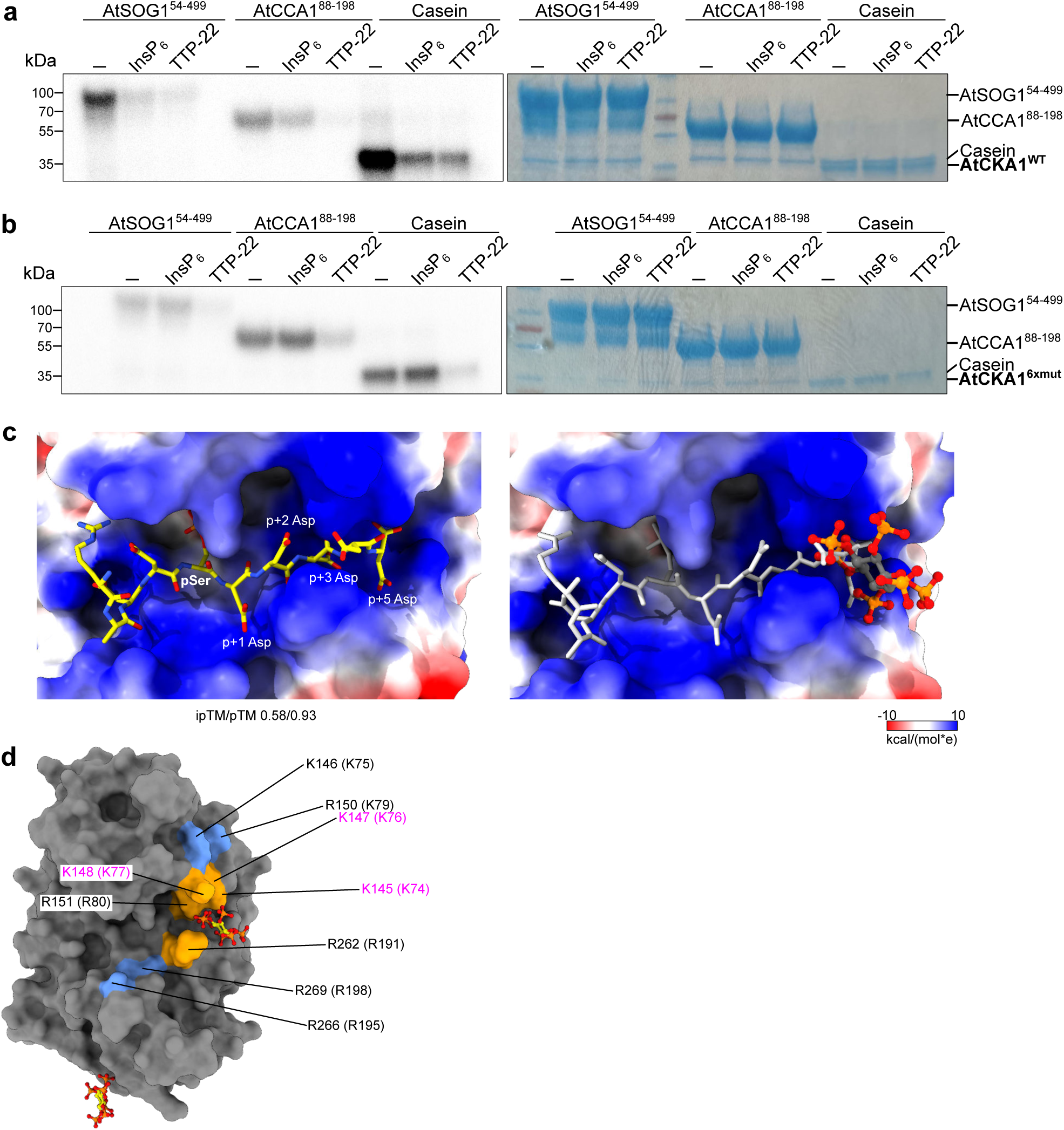
InsP6 reduces CK2 substrate phosphorylation by targeting the substrate binding surface. **a** In vitro transphosphorylation assay of AtCKA1 vs. AtSOG1, AtCCA1 or casein, and in the presence of either 200 μM InsP_6_ or 5 μM of the AtCKA inhibitor TTP-22. The autoradiograph is shown on the left, the corresponding Coomassie-stained SDS-PAGE gel on the right. **b** In vitro kinase assay for the AtCKA1^6xmut^ (K145A/K147A/K148A/K318A/H347A/K350A). **c** AlphaFold3 complex model of AtCKA1 bound to a generic CK2 peptide substrate (RRRAD<u>pS</u>DDDDD)^62^. Shown are electrostatic surface potentials in kcal/(mol*e) at 298 K from –10 (red) over 0 (white) to 10 (blue), mapped to the crystal structure of the AtCKA1 – InsP_6_ complex. The CK2 substrate peptide (left panel: in yellow, right panel: in gray) and InsP_6_ (in gray) are shown in bonds representation. **d** Molecular surface representation of the AtCKA1 (in gray) – InsP_6_ complex (in yellow, in bonds representation). Conserved basic residues involved in substrate binding and previously mutated in human CK2α^124^ are shown in orange if part of the InsP_6_ binding surface, otherwise in light blue. Labels refer to the amino-acid position in AtCKA1 (residues mutated in this study are highlighted in magenta), the corresponding residues in human CK2α (PDB-ID: pdb_00001jwh; r.m.s.d. with AtCKA1 is ∼1 Å comparing 331 corresponding C_α_ atoms)^59^ are shown in brackets.

While CK2 can phosphorylate diverse protein substrates, the presence of acidic amino acids at the positions p+1, p+2 and p+5 in synthetic substrate peptides strongly increases the phosphorylation efficiency of the enzyme^62^. To date, no structures of CK2 in complex with a protein substrate have been reported. We thus used the optimal synthetic substrate peptide RRRA-DD<u>S</u>DDDD (phosphorylation site underlined)^62,124^ to model a AtCKA1 enzyme – substrate complex with AlphaFold3 (Fig. 5c). In our model, the peptide is bound in the substrate binding cleft of AtCKA1 with the central phosphorylated Ser residue pointing towards the active site of the kinase. The aspartic acid residues in p+1, p+3 and p+5 are coordinated by basic amino acids in proximity of the binding cleft, which in part also form the binding site for InsP_6_ (Figs. 4e; 5c). Structural superposition of the AtCKA1 – peptide complex model and our AtCKA1 – InsP_6_ complex reveal severe steric clashes between InsP_6_ and the acidic C-terminus of the substrate peptide (Fig. 5c), rationalizing the reduced substrate phosphorylation activity of AtCKA1 in the presence of InsP_6_ (Fig. 5a).

A complete mutational analysis of conserved basic amino acids in human CK2 has been previously conducted^62,124^. Notably, replacing nine conserved basic residues with alanine greatly increased the Michaelis-Menten constant (K_M_) for synthetic substrate peptides, suggesting the presence of a basic substrate binding surface that coordinates the acidic residues in the substrates p-1 – p+5 position^62^. We mapped the corresponding amino acids onto the AtCKA1 – InsP_6_ complex structure and found that five of the nine residues are part of the proximal InsP _6_ binding site (shown in orange in Fig. 5d). This rationalizes the reduced trans-phosphorylation efficiency of AtCKA1^6xmut^ and further supports the idea that InsP_6_ negatively regulates AtCKA1 activity by occupying part of its positively charged substrate-binding surface (Fig. 5c,d).

### The InsP_6_/PP-InsP binding surfaces regulate CK2 function in vivo

We next sought to test if the InsP_6_/PP-InsP binding surfaces contribute to CK2 function in vivo. Several growth and developmental phenotypes have been previously reported for a *cka1/2/3* triple mutant, but no genetic complementation experiments were performed^77,78^. In our experiments, *cka1/2/3* mutants had primary roots similar to the wild type at seedling stage (Supplementary Fig. 5a,b), a reduced hypocotyl length in darkness (Supplementary Fig. 5c), and delayed flowering (Fig. 6a,b). We chose the most broadly expressed α-subunit AtCKA3, which shares 90% sequence identity at amino acid level with AtCKA1, for complementation assays (Supplementary Fig. 5d). Expression of *AtCKA3* from a constitutive promoter in the *cka1/2/3* background did not rescue the seedling phenotypes (Supplementary Fig. 5a-c). In contrast, *AtCKA3* could fully restore the flowering time of *cka1/2/3* mutants to wild type levels (Fig. 6a,b). Expression of a catalytically inactive version of the enzyme had no effect, whereas expression of the AtCKA3^6xmut^ InsP₆/PP-InsP binding surface mutant produced intermediate, variable flowering time phenotypes (Fig. 6a,b).

**Fig. 6:**
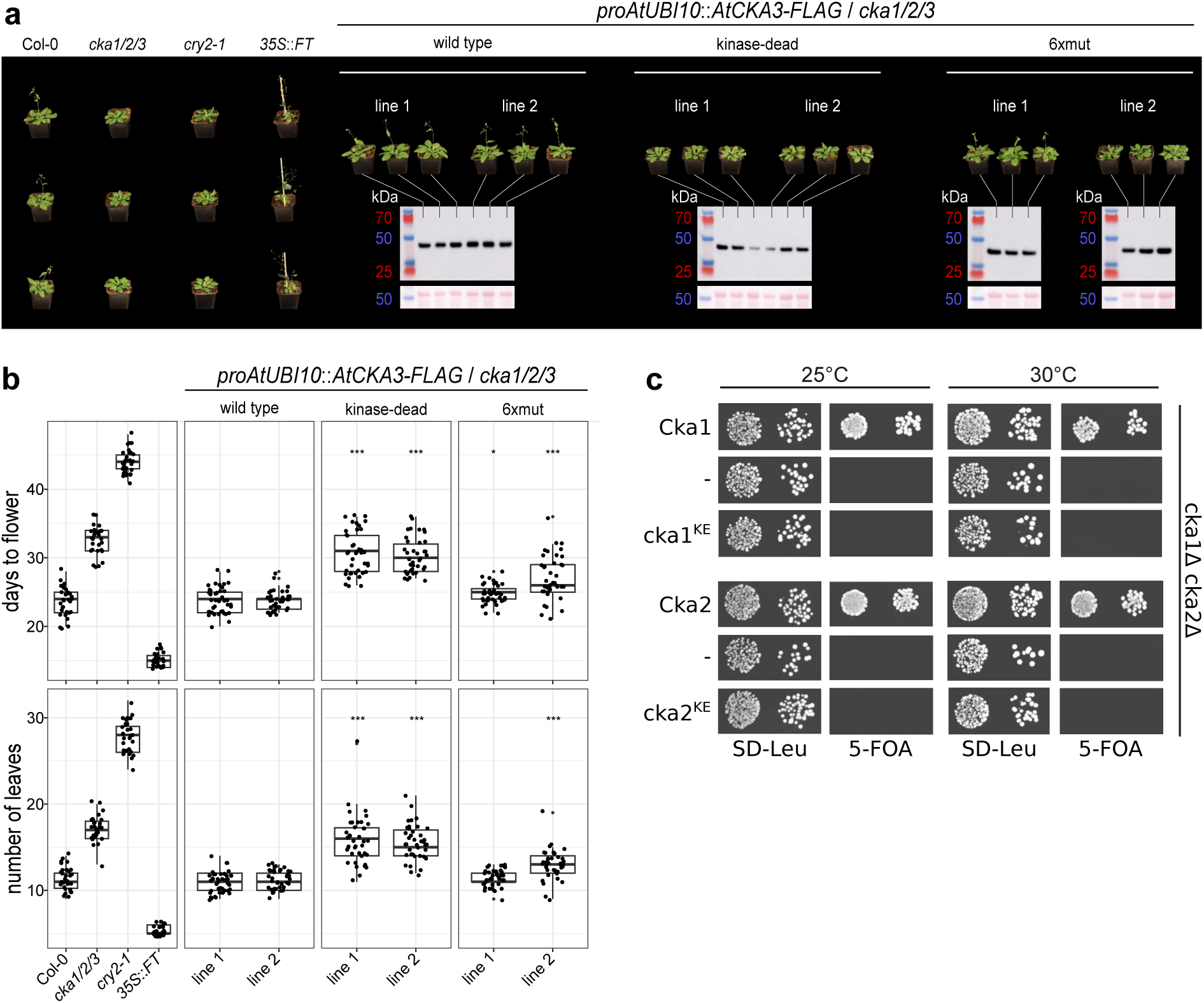
The InsP_6_/PP-InsP-binding surfaces contribute to CK2 function in Arabidopsis and in yeast. **a** Representative phenotypes 28 DAG (days after germination) for wild-type and mutant versions of AtCKA3 expressed in *cka1/2/3* background (pot size = 8×8 cm). Two independent transgenic T3 lines are shown for each construct together with the respective anti-FLAG western blot (theoretical MW of AtCKA3-3xFLAG is ∼42 kDa). **b** Box plots representing the days to flower (top) and number of leaves (bottom) of the lines shown in **a** (n=30-40). Bold black line, median; box, interquartile range (IQR); whiskers, lowest/highest data point within 1.5 IQR of the lower/upper quartile. Dunnett-type, two-sided multiplicity-adjusted p-values for the comparison against the Col-0 control are shown alongside (\**p* < 0.05; \*\*\**p* < 0.001). **c** Yeast cka1Δcka2Δ cells expressing either wild-type (Cka1, Cka2), empty vector (-) or InsP_6_/PP-InsP binding mutants (Cka1^KE^, Cka2^KE^) were spotted in a 10-fold dilution series on synthetic deficient (SD) media plates lacking leucine (Leu) and containing 5-Fluoroorotic acid (5-FOA), and incubated at the indicated temperatures.

The InsP_6_/PP-InsP binding surface is highly conserved among eukaryotic CK2 enzymes (Supplementary Fig. 3). In *Saccharomyces cerevisiae*, the kinase is encoded by two catalytic α-subunit isoforms, Cka1 and Cka2. Yeast strains lacking both isoforms (cka1Δ cka2Δ) are nonviable^125^. Expression of wild-type Cka1 or Cka2 fully complemented the lethality of the cka1Δ cka2Δ mutant (Fig. 6c). In contrast, mutant versions of Cka1 and Cka2, in which conserved basic residues within the InsP₆/PP-InsP binding surface adjacent to the nucleotide-binding cleft were substituted with glutamate, failed to rescue the cka1Δ cka2Δ mutant phenotype (Fig. 6c).

Collectively, these genetic experiments further support that the conserved InsP₆/PP-InsP binding surface contributes to CK2 function in vivo.

### InsP_6_/PP-InsP binding is independent from CK2 α_2_β_2_ holoenzyme formation

Arabidopsis CK2 exist either as an isolated α-subunit or as a heteromeric αβ complex^58^. We therefore investigated whether InsP_6_/PP-InsPs can regulate the formation or stoichiometry of the CK2 holoenzyme complex. The addition of InsP_6_ did not affect the reconstitution of an AtCKA1 – AtCKB1 complex in vitro (Fig. 7a). We then used size-exclusion chromatography coupled to multi-angle light scattering (SEC-MALS) to assess the oligomeric state of the purified AtCKA1 – AtCKB1 complex. We derived an apparent molecular weight of ∼135 kDa for the holoenzyme both in the presence and absence of InsP_6_, which is consistent with an α_2_β_2_ stoichiometry (Fig. 7b). Similarly, addition of InsP_6_ did not alter the monomeric state of the isolated AtCKA1 α-subunit (Fig. 7b).

**Fig. 7:**
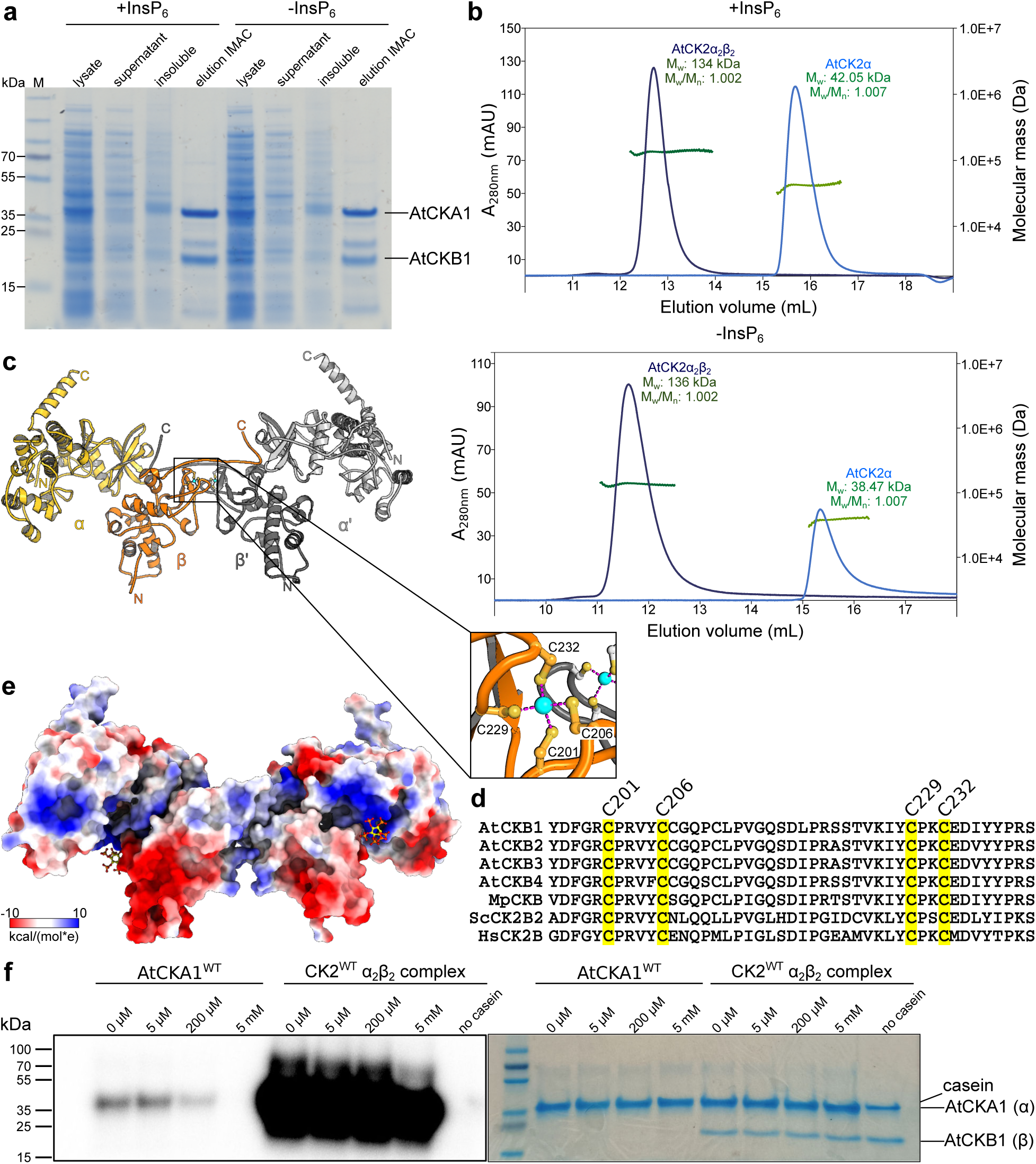
The AtCKA1 – AtCKB1 α_2_β_2_ holoenzyme can associate with InsP_6_ but is not regulated by it. **a** SDS-PAGE analysis of an Immobilized Metal Affinity Chromatography (IMAC) performed in the presence or absence of 1 mM InsP_6_. Untagged AtCKB1 (theoretical MW ∼22.2 kDa) co-purifies with 6xHis-tagged AtCKA1 (MW ∼40.3 kDa), both recombinantly expressed in *E. coli*. **b** SEC-MALS profiles of the AtCKA1 – AtCKB1 complex (dark blue/ dark green green) and the isolated AtCKA1 α-subunit (light blue / light green) in the presence (top) or absence (bottom) of 1 mM InsP_6_. The theoretical MW of the α_2_β_2_ heterotetramer is ∼125 kDa. **c** Crystal structure of the AtCKA1 – AtCKB1 complex. Shown are ribbon diagrams of the α and α1 (in yellow, and light gray), as well as the β and β1 (in orange and dark gray, respectively) subunits. Two Zn^2+^ ions are highlighted as magenta spheres. The inset provides details of the AtCKB1 Zn ^2+^ binding site. Cys residues are highlighted in bonds representation, ion coordination is depicted by dotted lines (in magenta). **d** Multiple sequence alignment of the Arabidopsis (At), Marchantia (Mp), yeast (Sc) and human (Hs) β-subunits, with the Cys residues contributing to the structural Zn ^2+^ binding motif highlighted in yellow. **e** Electrostatic surface potential in kcal/(mol*e) at 298 K from –10 (red) over 0 (white) to 10 (blue) mapped to the crystal structure of AtCKA1 – AtCKB1 complex. Bound InsP _6_ molecules are highlighted in bonds representation. **f** In vitro transphosphorylation assay of AtCKA1 vs the AtCKA1 – AtCKB1 α_2_β_2_, holoenzyme in the presence of either 0, 5 μM, 200 μM, or 5 mM InsP_6_ and using casein as substrate. The autoradiograph is shown on the left, the corresponding Coomassie-stained SDS-PAGE gel on the right.

We obtained crystals of the AtCKA1 – AtCKB1 complex in the presence of 5 mM InsP_6_ and determined the low-resolution structure of the α_2_β_2_ holoenzyme (Supplementary Table 3). In our complex structure, the β-subunits are stabilized by conserved structural Zn^2+^-binding motifs (Fig. 7c,d)^126^ and form the center of a butterfly-shaped assembly, as previously described for the human holoenzyme (Fig. 7c; Supplementary Fig. 6)^59^. Two strong difference map peaks in the vicinity of the nucleotide-binding cleft were modeled as InsP_6_ molecules (Fig. 7e). InsP_6_ is bound by the conserved basic surface patch in CK2α, which, in the complex structure, is in close proximity to a highly acidic surface patch within the β-subunit (Fig. 7e). Trans-phosphorylation assays comparing the monomeric AtCKA1 with the AtCKA1 – AtCKB1 complex revealed that the holoenzyme’s catalytic activity is drastically higher than that of the isolated α-subunit (Fig. 7f). The addition of increasing concentrations of InsP_6_ inhibited the isolated α-subunit, but not the AtCKA1 – AtCKB1 complex (Fig. 7f). Overall, our experiments suggest that, although InsP_6_/PP-InsPs can bind both the isolated CK2 α-subunit and the α_2_β_2_ holoenzyme, these metabolites regulate the trans-phosphorylation activity of CK2 when present in its monomeric α-subunit state.

### Plant CK2 α-subunits can function independently from their regulatory β-subunits

This prompted us to investigate whether CK2 in Arabidopsis functions as a hetero-tetrameric holoenzyme or as a free α-subunit, or in both configurations. No loss-of-function phenotypes have been reported for the four CK2 β-subunits present in Arabidopsis. We therefore used CRISPR/Cas9 gene editing to generate a *ckb1 ckb2 ckb3 ckb4* (*ckb1/2/3/4*) quadruple mutant, as well as a single *cka4* knock-out mutant for the chloroplast-localized α-subunit (Supplementary Fig. 7). We found that *ckb1/2/3/4* seedlings had shorter primary roots than the Col-0 wild type (Fig. 8a). The *cka1/2/3* mutant behaved similar to wild type (also see Supplementary Fig. 5a,b), whereas primary root growth was reduced in the *cka4* mutant, as previously reported (Fig. 8a)^80^. The shoots of the *ckb1/2/3/4* and *cka4* mutants had higher levels of anthocyanins than the wild-type control seedlings (Fig. 8b). In the dark, the hypocotyl length was significantly reduced in the *cka1/2/3* and *ckb1/2/3/4* mutants when compared to the wild type, the *cka4* mutant exhibited no visible phenotype under these conditions (Fig. 8c).

**Fig. 8:**
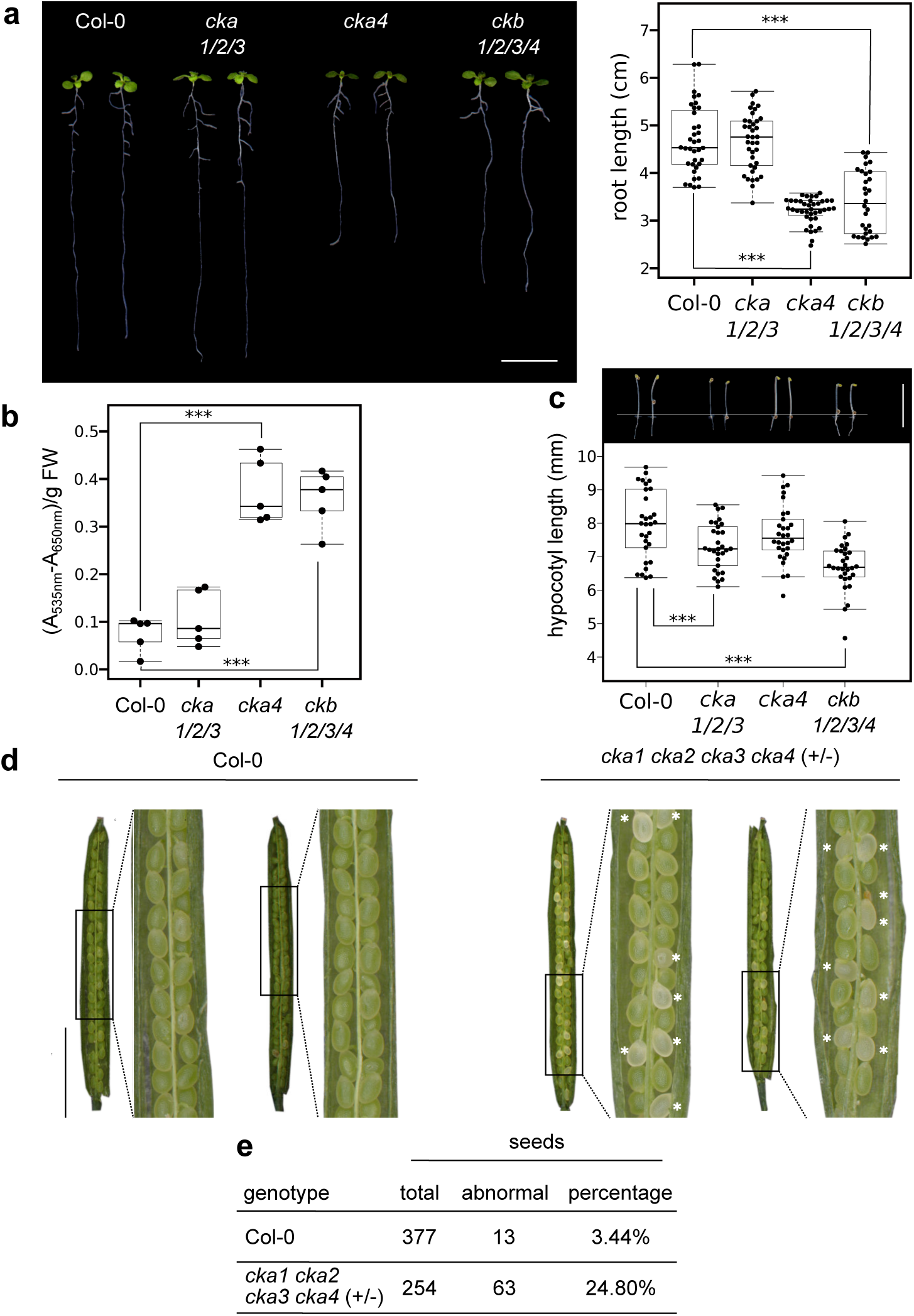
Arabidopsis CK2 α-subunits can function independently from their regulatory β-subunits. **a** Primary root length phenotypes of *cka1/2/3*, *cka4* and *ckb1/2/3/4* mutant vs. the Col-0 wild-type control. Seeds were stratified and left to germinate for 10 d. Representative seedlings from each line are shown on the left (scale bar = 1cm), quantifications (n=30) are shown on the right. Center black line, median; box, interquartile range (IQR); whiskers, lowest/highest data point within 1.5 IQR of the lower/upper quartile. Dunnett-type, two-sided multiplicity-adjusted *p*-values for the comparison against the Col-0 control are shown alongside (\*\*\**p* < 0.001). **b** Shoot anthocyanin content of 10 d-old seedlings (n=5). **c** Hypocotyl length phenotypes of seedlings grown for 3 d in the dark. Representative seedlings are shown on top (scale bar = 1cm), quantifications are shown below (n=30). **d** Aborted seed phenotype of the *cka1/2/3/4* quadruple mutant. Shown are representative dissected siliques, Col-0 (n=6) is shown on the left, the *cka1 cka2 cka3 cka4* (-/+) (n=4) is shown on the right (scale bar = 0.5 cm). The insets provide a zoomed-in view, white asterisks highlighting aborted white seeds. **e** Quantification of total vs. aborted seeds in the different genotypes.

Because the *cka1/2/3* triple mutant displayed comparatively mild phenotypes relative to our *ckb1/2/3/4* mutant, we next examined the genetic relationship between the nuclear-localized α-subunits AtCKA1-3 and the chloroplast-localized AtCKA4. Using CRISPR/Cas9, we obtained several plants with CRISPR-events targeting *CKA4* in the *cka1/2/3* background (Fig. 8d). Siliques from segregating *cka1 cka2 cka3 cka4* (+/-) lines exhibited a markedly increased frequency of aborted seeds and showed distorted segregation patterns, indicating that simultaneous loss of all four α-subunits is embryo lethal (Fig. 8d,e). This severe phenotype of the *cka* quadruple mutant, contrasted with the mild growth defects of the *ckb* quadruple mutant, strongly suggests that CK2 α-subunits can function independently of the regulatory β-subunits in Arabidopsis.

To extend our observations beyond Arabidopsis, we generated CRISPR/Cas9 knockout lines for the single *MpCKA* and *MpCKB* genes in *Marchantia polymorpha* (Supplementary Fig. 7). The *cka^ge^*mutant exhibited a pronounced growth defect, displaying a markedly reduced thallus area relative to the Tak-1 control (Fig. 9a,b). In contrast, the *ckb^ge^* mutant showed growth dynamics comparable to Tak-1 (Fig. 9a,b). As older *cka^ge^*thalli visually accumulated pigments, we quantified UV-B-absorbing metabolites, as previously described^127^. Consistent with their pigmentation phenotype, *cka^ge^* but not *ckb^ge^*mutants showed elevated levels of UV-B-absorbing compounds, possibly flavones (Fig. 9c). We attempted genetic complementation of *cka^ge^* mutant with gRNA resistant versions of MpCKA^wild type^, MpCKA^K157M/D258N^ (kinase dead) of MpCKA^K157A/K159A/K160A/K330A/H359A/K362A^ (6xmut), each expressed from the constitutive *proMpEF1* promoter and fused to a C-terminal RFP tag. However, no transformants were recovered in the *cka^ge^*background. As an alternative strategy, we expressed these constructs in Tak-1 (Fig. 9d). The InsP_6_/PP-InsP binding mutant did not differ from wild type in thallus growth or pigment accumulation (Fig. 9a-c). In contrast, expression of the kinase-dead *MpCKA* variant resulted in reduced thallus size and increased accumulation of UV-B-absorbing compounds (Fig. 9a-c). This dominant-negative effect is potentially caused by the catalytically inactive MpCKA interacting with CK2 substrates, titrating them away from the active α-subunit present in Tak-1.

**Fig. 9:**
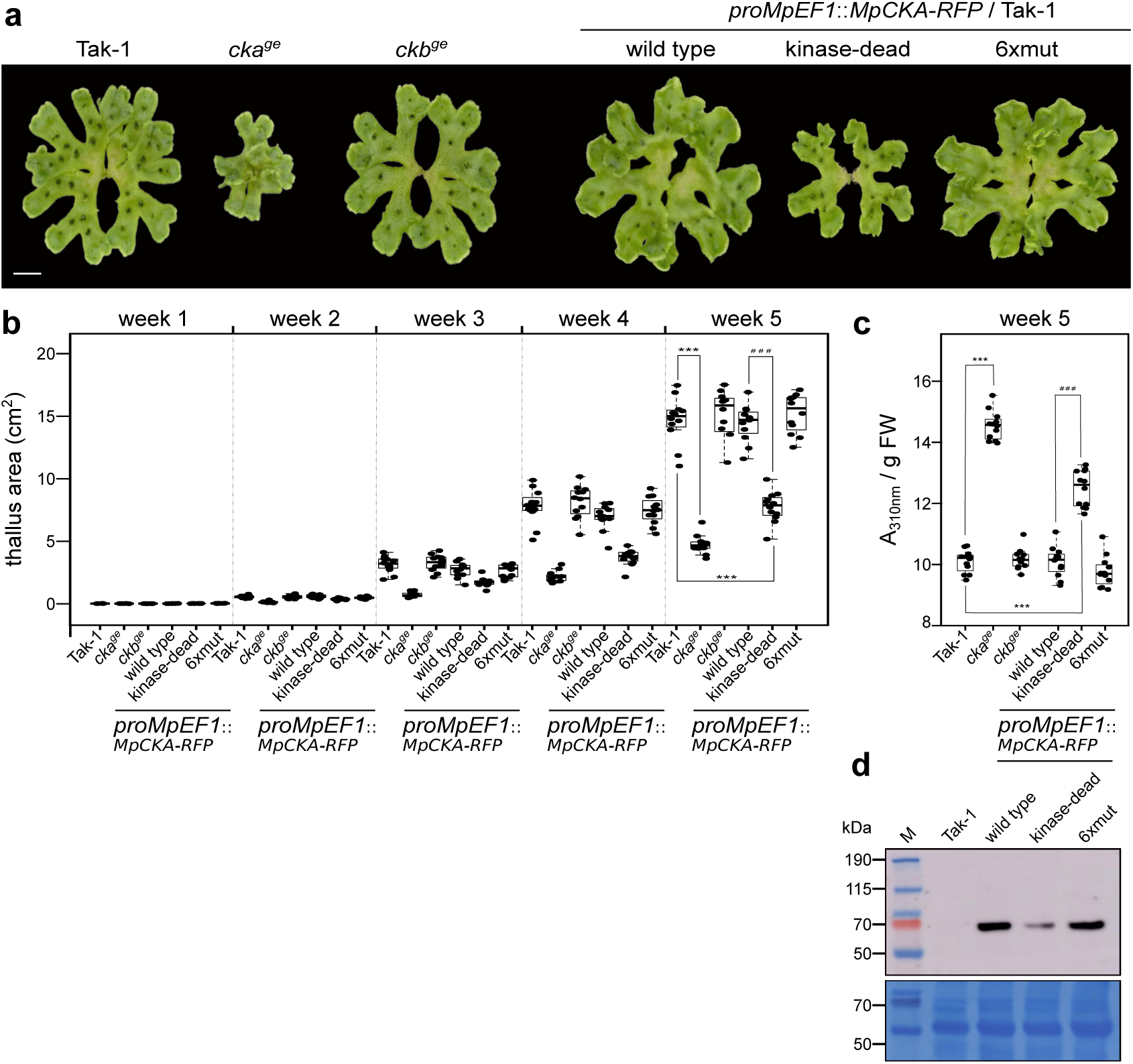
Deletion of of the single *MpCKA* but not the *MpCKB* gene strongly affects Marchantia growth and development. **a** Representative phenotypes of 5 week-old plants (scale bar = 1 cm; kinase-dead, MpCKA^K157M/D258N^; 6xmut, MpCKA^K157A/K159A/K160A/K330A/H359A/K362A^). **b** Thallus surface areas of the genotypes shown in **a** in a time course experiment (n=12). Statistical significance was assessed at week 5 using a two-sided Dunnett test with the Tak-1 wild type (\*\*\**p* < 0.001), or the *proMpEF1*::*MpCKA-RFP* (^###^*p* < 0.001) as reference. **c** UV-B-absorbing compounds content quantified at week 5 (n=12). **d** Western blot analysis of *proMpEF1*::*MpCKA-RFP* overexpressing lines probed with a pan-RFP polyclonal antibody.

Taken together, our genetic analyses in Arabidopsis and *Marchantia* indicate that CK2 α-subunits play essential roles in plant growth and development, whereas β-subunits are largely dispensable under laboratory conditions. These results suggest that the α-subunit can function independently of the canonical α₂β₂ holoenzyme in planta, and therefore its modulation by InsP₆/PP-InsP is likely of physiological relevance.

## Discussion

### InsP_6_/PP-InsP interactomes define novel and conserved PP-InsP receptors

Inositol pyrophosphates have emerged as important nutrient messengers in plants^2,5,11,20,22,25,26^, and their mechanistic contributions to Pi homeostasis^11,26,39,40,46,47^ and auxin responses^6,34^ are relatively well characterized. Here we present a system-wide interaction screen, which suggests the presence of many additional, currently uncharacterized InsP_6_/PP-InsP receptors in Arabidopsis. The strong overlap between our InsP_6_, InsP_7_ and InsP_8_ samples as well as within the human data set^92^ suggests that many putative interactors cannot efficiently discriminate between PP-InsP isomers, at least at the biochemical level (Fig. 1a). Furthermore, the absence of known PP-InsP interacting proteins present in seedlings such as AtVIH1/2^17^, AtPHO1^49^, AtNLA^128^, AtTIR1^6,34^ or AtCOI1^9,35,37^ suggest that our dataset is incomplete. The significant overlap between our interactome and InsP_6_-bound complex structures in the PDB strongly argues for a conserved function for InsP_6_ or PP-InsPs in the assembly and function of conserved molecular machines such as the spliceosome (Fig. 2d), the ribosome (Fig. 2f; 3d) and the polyadenylation machinery (Fig. 3c). This rationalizes at least in part the abundance of spliceosomal, ribosomal and nucleolar proteins in the fungal, human and plant PP-InsP interactomes^91–93^ (Fig. 1). Our structure-guided validation pipeline (Fig. 2g) allowed us to filter many putative InsP_6_/PP-InsP interactors into a consistent set of putative direct binders and binder-associated complex subunits, which together cover a surprisingly large fraction of the total interactome (Figs. 2;3; Supplementary Table 1). Future studies will clarify the roles of PP-InsPs in mRNA splicing, polyadenylation, ribosome maturation/function, and in cell signaling.

### CK2 α-subunits are conserved eukaryotic InsP_6_/PP-InsP receptors

The lack of PP-InsP isoform selectivity in our screen prompted us to perform an orthogonal interaction screen with the PPIP5K AtVIH2, which generates the nutrient messenger InsP_8_^8,9,11,17^. This approach has been recently used to identify a bona fide PP-InsP receptor in *Marchantia polymorpha*^27^. The substantial overlap between the InsP_6_/PP-InsP and the AtVIH2 interactomes suggests that the incorporation of a given PP-InsP isomer into a binding protein is influenced not only by that protein’s intrinsic binding affinity and kinetic properties, but also by its spatial proximity to a PP-InsP-synthesizing enzyme (Fig. 4a-c).

All nuclear-localized α-subunits of CK2 were recovered as putative InsP_6_/PP-InsP binders from our screen (Supplementary Fig. 1; Supplementary Table 1), and AtCKA1 was also identified and validated as a AtVIH2 interactor (Fig. 4a-c). Importantly, the α-subunits of yeast and human CK2 form part of their respective PP-InsP interactomes^92,93^, and human CK2 has been shown to directly bind InsP_6_ and InsP_7_, respectively^54,55^. The AtCKA1 – InsP_6_ complex structure and kinetic binding assays together suggest that plant, fungal and human CK2 share conserved InsP_6_/PP-InsP binding sites (Fig. 4e,f; Supplementary Fig. 3)^54,55^. The plant enzyme does not inherently discriminate between InsP_6_, InsP_7_, or InsP_8_; however, its association with AtVIH2 suggests that InsP_8_ may represent the physiologically relevant ligand (Fig. 4a-c). Importantly, we could demonstrate that InsP_6_ competes with CK2 substrates for binding to the basic substrate binding patch in the CK2 α-subunit (Fig. 5). This suggests that phosphorylation of target proteins with acidic CK2 substrate-binding motifs may be regulated by InsP_6_/PP-InsPs^52,53^, the concentration of which fluctuates in response to changing nutrient conditions and during the cell cycle^5,129^. Mutations within the InsP_6_/PP-InsP binding site revealed that the surface area contributes substantially to Cka1 and Cka2 function in yeast (Fig. 6c). In contrast, mutations in the corresponding region of AtCKA3 (AtCKA3^6xmut^) had only a modest effect on flowering time regulation in Arabidopsis (Fig. 6a,b). Nevertheless, InsP_6_ clearly inhibited the ability of AtCKA1 to phosphorylate the known flowering time regulator AtCCA1 in vitro (Fig. 5a)^81,84,85^. Given the large number of potential CK2 substrates^130,131^ it is difficult to pinpoint physiologically relevant CK2 – substrate interactions regulated by InsP_6_/PP-InsP in planta. Notably, CK2 has been mechanistically linked to phosphate homeostasis in Arabidopsis and in rice^79,89,132,133^, a process that is centrally regulated by InsP_8_^38^.

### The activity of isolated α-subunits is regulated by InsP_6_/PP-InsPs

CK2 has been characterized as a constitutively active, pleotropic protein kinase^60,134^. Arabidopsis has four catalytic α-subunits and four regulatory β-subunits, whereas in Marchantia both isoforms are encoded by single-copy genes. Our reconstituted AtCKA1 – AtCKB1 complex forms the canonical α_2_β_2_ holoenzyme and is structurally related to the human CK2 complex (Fig. 7c)^59^. The presence of the β-subunit strongly stimulates the trans-phosphorylation activity of CK2 (Fig. 7f), as previously reported^58^. While InsP_6_ efficiently inhibited the ability of isolated AtCKA1 to transphosphorylate different substrates, no such effect was observed for the AtCKA1 – AtCKB1 complex (Fig. 7f), suggesting that InsP_6_/PP-InsP are regulators of the isolated CK2 catalytic module, not of the α_2_β_2_ holoenzyme. This led us to investigate the relative contributions of the α-and β-subunits to the growth and development of Arabidopsis and Marchantia. Our genetic experiments suggest essential functions for the catalytic α-subunits in both Arabidopsis and Marchantia, while knock-out mutants of the β-subunits display rather mild phenotypes in both organisms (Figs. 8;9). This suggests that CK2 can operate as a catalytic monomer independently from its regulatory β-subunit in plants and may be regulated by other mechanisms including InsP_6_/PP-InsPs under these circumstances. The conservation of the InsP_6_/PP-InsP binding sites in the CK2 α-subunit suggests that this may represent a conserved regulatory mechanism in fungi, plants, and animals.

### Does the InsP_6_/PP-InsP binding site in CK2 contribute to protein pyrophosphorylation?

InsP₇-mediated protein pyrophosphorylation was originally identified in vitro, in yeast and in human cells^68,69^. This modification depends on a priming phosphorylation event catalyzed by CK2^69^. The animal InsP_6_ kinase IP6K1 interacts with both CK2 and with pyrophosphorylated target proteins^67^. We have recently reported that protein pyrophosphorylation is widespread in human cells^70^ and a physiological function for protein pyrophosphorylation in plant phosphate homeostasis, photomorphogenesis and jasmonic acid signaling has been proposed^135^. Our experiments suggest that plant CK2 α-subunits can interact with both PP-InsPs and the PPIP5K AtVIH2 (Fig. 4a-c, f). This indicates that CK2 may act as a priming kinase for protein pyrophosphorylation as well as a scaffold that brings PP-InsP donors and protein substrates into close proximity. Formation of an AtCK2 – AtVIH2 complex could therefore coordinate the kinase and pyrophosphate donor activities required for protein pyrophosphorylation. Our *Arabidopsis* and Marchantia PPIP5K and CK2A knockout lines^11,20^ (Figs. 8;9) will facilitate the testing of this hypothesis at a systems level and may contribute to the mechanistic analysis of inositol pyrophosphate-dependent protein pyrophosphorylation in plants.

## Materials and methods

### InsP_6_/PP-InsP interaction screen

#### Plant material

For the PP-InsP interaction screen, *Arabidopsis thaliana* Col-0 seedlings were grown on ^½^Murashige and Skoog (^½^MS, Duchefa, 1% (w/v) sucrose, 1 mM K_2_HPO_4_/KH_2_PO_4_) 0.8% (w/v) agar plates pH 5.7 under long-day conditions (16 h light / 8 h dark cycle) at 22°C for one week. They were then transferred to Pi starvation ^½^MS agar (^½^MS, 1% (w/v) sucrose, 20 μM KH_2_PO_4_/K_2_HPO_4_, 980 μM KCl). After one week of Pi starvation, seedlings were harvested, shock frozen in liquid nitrogen and stored at –80°C until further use.

#### Pulldown screen

For protein extraction, protein samples were homogenized using a MM400 (Retsch) and incubated with buffer (50 mM HEPES pH 7.5, 150 mM NaCl, 1 mM EDTA, 10% [v/v] glycerol, 0.1% [v/v] Triton X-100, 1 mM DTT) including phosphostop (Roche) and plant protease inhibitor mix (Merck Millipore) for 1 h at 4°C with gentle rotation. Samples were then centrifuged at 15,000 × g for 10 min at 4°C. The centrifugation step was repeated with the supernatant. Protein concentrations was measured using a Bradford assay^136^ and adjusted to 1 mg/mL with buffer. To prepare the PP-InsP coupled beads, lyophilized 1/3-biotin-InsP_6_, 1/3-biotin-5PCP-InsP_5_ or 3-biotin-1,5-(PCP)_2_-InsP_4_^91,92^ were solubilized in H_2_O. For each reaction, 20 nmol of the respective PP-InsP was immobilized on streptavidin-coupled Sepharose beads (Sigma) equilibrated in PBS for 1h with gentle shaking. The beads were then washed once with PBS and equilibrated in buffer. Per reaction, 150 µL of purified protein extraction lysate was added to the PP-InsP beads for 3h at 4°C, the total concentration of the final probe was 150 µM. Reactions were washed with 25 mM HEPES pH 7.5, 150 mM NaCl, 1 mM EDTA in a 96-well plate using a vacuum manifold. 5 mM stock solutions of soluble InsP _6_, 5PCP-InsP_5_ and 1,5(PCP)_2_-InsP_4_^90^ were prepared in 25 mM HEPES pH 7.5, 150 mM NaCl, 1 mM EDTA. The enriched proteins were eluted by incubating the beads with 50 µL of the corresponding soluble PP-InsP at 4°C with shaking. This step was repeated once, resulting in 100 µL of eluate per reaction, which was subsequently shock frozen in liquid N_2_ and stored at –80°C.

#### Mass spectrometry analysis

Samples were dried under vacuum and further processed using the iST kit (Preomics) following the manufacturer’s instructions. Briefly, samples were resuspended, followed by denaturation, reduction and alkylation of the samples. Digestion was performed with a trypsin/LysC before the samples were dried again under speed vacuum. For analysis, samples were dissolved in 5% (v/v) CH_3_CN, 0.1% (v/v) HCOOH and injected into a Q-Exactive F hybrid quadrupole Orbitrap mass spectrometer (Thermo Fisher) equipped with an Easy nLC100 liquid chromatography system. Samples were separated for 90 min on a C18 column using an H_2_O to CH_3_CN gradient containing 0.1% (v/v) HCOOH. MS2 analysis was performed using dynamic exclusion. Raw data were processed using Proteome Discoverer 2.4 (Thermo Fisher). Extracted spectra were searched against the *A. thaliana* reference proteome for hit identification. For label-free MS1 quantification, peaks were detected using the Minora Feature Detector node. The false discovery rate (FDR) was set at 1 % and at least two unique peptides per protein were used.

#### Bioinformatic analysis

The degree of enrichment was calculated for each sample (InsP_6_, InsP_7_, InsP_8_) and as an average for all PP-InsP samples. Enrichment values greater than 1.5-fold were classified as hits. Gene ontology (GO) analyses were performed in R/Bioconductor using the topGO package^137,138^. Comparative interactome analyses were conducted between plant, yeast, and human InsP_7_ datasets. Local BLAST databases were generated using 526 hits from the *A. thaliana* (1/3-biotin-5PCP-InsP_5_ sample; Supplementary Table 1), 82 hits from *S. cerevisiae* (+EDTA sample)^93^ and 278 hits from *H. sapiens* HEK293T cells (tri-InsP_7_ sample) ^91^. Blast searches were performed against these local databases with blastp^139^ with an e-value cutoff of 1e−5, and only unique top hits were retained for subsequent analyses. InsP_6_-bound structures were retrieved from the Protein Data Bank (PDB; http://www.rcsb.org, September 2025) using the chemical identifier IHP. The amino acid sequences of all unique chains from 304 IHP-containing structures were compared to a local BLAST database of 595 putative InsP_6_/PP-InsP interactors using blastp (e-value cutoff of 1e-5)^139^. The number of hits per PDB entry and unique chain identifier was mapped. Using this approach, 47 unique interactome proteins were associated with 61 subunits of the human pre-catalytic spliceosome precursor complex (PDB ID: pdb_00006ah0)^97^, considering only the top hits with at least 75% sequence coverage. For remaining complexes, AlphaFold2 models ^106^ were obtained for each unique *Arabidopsis* sequence corresponding to a PDB chain ID. Structural superpositions between AlphaFold2 models and experimental structures were performed in ChimeraX (v1.10.1) using the Matchmaker tool^140^. Missing structures and protein complex models were generated using AlphaFold3^111^ pTM and ipTM values are reported in the respective figure panels. InsP_6_ docking (chemical ID IHP) was performed with Boltz-2 via the protein – ligand co-folding workflow (RowanSci, https://rowansci.com/); predicted IC_50_ values are reported in the respective figure legend. Structural representations were done in ChimeraX (version 1.10.1)^140^.

### Immunoprecipitation mass-spectrometry

Seedlings of *promAtVIH2*::*AtVIH2-mCit* (NASC ID: N2109981) in the *vih1-2* (SALK_094780C) *vih2-4* (GK_080A07) double mutant background^11^, along with a *35S*::*YFP* control line in the Col-0 ecotype, were grown for 14 d on ^½^MS medium (Duchefa). The immunoprecipitation followed by mass-spectrometry (IP/MS) experiment was performed as previously described^141^. Briefly, plant tissues were ground in liquid N_2_ and homogenized in lysis buffer containing 50 mM Tris-HCl (pH 7.5), 150 mM NaCl, 2 mM EDTA, 1% (v/v) Triton X-100, cOmplete™ EDTA-free protease inhibitor cocktail (Roche), and PhosSTOP™ phosphatase inhibitor (Roche). After solubilization on ice for 30 min, cell debris was removed by two rounds of centrifugation at 14,000 × g for 15 min at 4°C. The resulting supernatants were incubated with 50 μl GFP-Trap magnetic agarose beads (ChromoTek) for 3 h at 4°C to allow protein binding. The beads were then washed four times with wash buffer (50 mM Tris-HCl pH 7.5, 150 mM NaCl, 1 mM EDTA) containing 0.1% (v/v) Triton X-100, followed by one wash without Triton X-100. On-bead protein digestion was carried out using a trypsin/LysC mix in combination with iST kits (PreOmics). The resulting peptides were analyzed by nanoLC-ESI-MS/MS using an Easy-nLC 1000 system (Thermo Fisher Scientific) coupled to a Q-Exactive HF Hybrid Quadrupole-Orbitrap mass spectrometer. Peptide identification was performed using Mascot (Matrix Science) against the *Arabidopsis thaliana* reference proteome database (ToxoDB) along with the bait protein sequences. Data processing and validation were conducted using Proteome Discoverer (Thermo Fisher Scientific), applying a 1% false discovery rate (FDR) threshold at the protein level and requiring a minimum of two unique peptides per protein. 472 hits with a log_2_ ratio ≥ 0.58 and a *p*-value ≤ 0.05 were compared to a local blast database containing 595 putative InsP_6_/PP-InsP interacting proteins.

### Split luciferase assay

The split-luciferase complementation assay was performed as previously described^142^. Full-length coding sequences of AtVIH2, AtCKA1, and AtCKA4 were cloned into the pGWB-Cluc and pGWB-Nluc vectors, respectively, using the Gateway LR recombination system (Invitrogen). The pCAMBIA1300-Cluc vector was used as a negative control^143^. Constructs were introduced into *Agrobacterium tumefaciens* strain GV3101, and positive colonies were cultured for 16 h at 30°C before being resuspended in induction buffer (10 mM MES, pH 5.7, 10 mM MgCl₂, and 150 mM acetosyringone) to an OD_600 nm_ of 1.0. The agrobacterial suspensions were infiltrated into the abaxial side of leaves from 4-week-old *Nicotiana benthamiana* plants. Following a 48 h incubation in a growth chamber, luciferase activity was detected by applying 1 mM luciferin potassium salt solution (Invitrogen), and luminescence was measured using the NightShade imaging system (LB985, Berthold Technologies).

### Protein expression and purification

AtCKA1 (residues 76-409, Uniprot-ID Q08467, http://uniprot.org), AtCKA3 (residues 1-331, A0A178VTA8) were cloned from a seedling cDNA library into plasmid pMH-HC providing a non-cleavable C-terminal 6xHis tag. AtCCA1 (residues 88-198, A0A178VLU6) and AtSOG1 (residues 54-499, A0A178WLI1) were cloned from synthetic genes (Twist Biosciences) into plasmid pMH-HsmbpT providing a 6xHis-tagged, tobacco etch virus (TEV)-cleavable maltose binding protein fusion tag. AtCKB1 (residues 95-287, A0A5S9YCV4) was cloned from a synthetic gene (Twist Biosciences) into plasmid pMH-HC with a stop codon 5’ of the C-terminal 6xHis tag. Protein expression in *E. coli* Rosetta 2 (DE3) cells grown to OD_600 nm_= 0.6 was induced with 0.3 mM ispropyl β-D-galactoside in terrific broth at 18°C for 16 h. Cells were collected by centrifugation at 4,000 × g for 20 min at 4°C, washed in PBS buffer, centrifuged again at 4,000 × g for 15 min, resuspended in 50 mM HEPES (pH 7.5), 500 mM NaCl, 1 mM MgCl_2_, 0.1% (v/v) Triton X-100 in 1:5 (g/mL) ratio, snap-frozen in liquid N_2_, and stored at –20°C until further use.

For protein purification of AtCKA1 and AtCKA3, cells pellets from 6L of culture were thawed, supplemented with DNAse I (Thermo Fisher Scientific), lysozyme and cOmplete EDTA-free protease inhibitor (Merck), imidazole (pH 8.0) to a final concentration of 20 mM, and homogenized. Cells were lyzed by sonication (Branson DS450) and the lysate was cleared by centrifugation at 45,000 × g for 30 min at 4°C and filtrated using a 0.45 µm molecular weight cut-off (MWCO) filter (Pall Corporation). The supernatant was loaded onto a Ni^2+^ affinity column (HisTrap HP 5ml, Cytiva), equilibrated in buffer (20 mM HEPES [pH 7.5], 300 mM NaCl). The column was washed with 5 column volumes (CV) of buffer a supplemented with 50 mM imidazole (pH 8.0), and eluted in buffer A supplemented with 500 mM imidazole (pH 8.0). The eluted protein fractions were analyzed by SDS-PAGE, pooled, concentrated to 5 mL and loaded onto a HiLoad Superdex 200 pg 16/600 colum (Cytvia) equilibrated in 20 mM HEPES (pH 7.5), 150 mM NaCl at a flow rate of 0.75 mL/min. Monomeric peak fractions were pooled and concentrated with an Amicon centrifugal filter (Merck, 10 kDa MWCO) to 5-15 mg/mL. Point mutations were introduced by site-directed mutagenesis and the different variants (AtCKA1^N-lobe^, K145A/K147A/K148A; AtCKA1^C-lobe^, K318A/H347A/K350A; AtCKA1^6xmut^ K145A/K147A/K148A/K318A/H347A/K350A) were purified using the same protocol as described for the wild type AtCKA1. AtCCA1 was purified using the same purification scheme. AtSOG1 was purified by Co^2+^ metal affinity chromatography in batch by mixing the filtered supernatant with 4 mL of TALON resin (Takara Bio) equilibrated in 20 mM HEPES (pH 7.5), 300 mM NaCl for 1 h at 4°C. The beads were washed in the same buffer supplemented with 5 mM imidazole (pH 8.0), and batch elution was done in 20 mM HEPES (pH 7.5), 150 mM NaCl, 250 mM imidazole. The N-terminal MBP fusion tag was left intact and the MBP-AtSOG1 and MBP-AtCCA1 fusion proteins were further purified by size-exclusion chromatography on a HiLoad Superdex 200 pg 16/600 colum (Cytvia) equilibrated in 20 mM HEPES (pH 7.5), 150 mM NaCl.

To purify the AtCKA1 – AtCKB1 complex, AtCKA1 with an C-terminal 6xHis tag and untagged AtCKB1 were expressed separately in *E. coli* as described above. The resulting cell pellets were mixed in 1:1 ratio, homogenized and lyzed together. The lysate was cleared by centrifugation at 45,000 × g for 30 min at 4°C, filtrated using a 0.45 µm molecular weight cut-off (MWCO) filter (Pall Corporation) and supplemented with imidazole (pH 8.0) to a final concentration of 20 mM. The supernatant was loaded onto a 5 mL HisTrap HP column (Cytiva) equilibrated in buffer (20 mM HEPES [pH 7.5], 300 mM NaCl). The column was then washed with 5 CVs of buffer supplemented with 50 mM imidazole (pH 8.0). Protein elution was done by applying an imidazole gradient from 50 mM to 500 mM over 10 CV. Peak fractions were analyzed by SDS-PAGE and fractions containing the AtCKA1 – AtCKB1 complex were pooled, concentrated to 5 mL (Amicon centrifugal filter, 30 kDa MWCO, Merck) and loaded onto a HiLoad Superdex 200 pg 16/600 (Cytiva) equilibrated in 20 mM HEPES pH 7.5, 300 mM NaCl to separate the heterotetrameric complex from individual AtCKA1 α-subunits. All purified samples were concentrated to 5-15 mg/mL using Amicon centrifugal filters (Merck, 10-30 kDa MWCO) and snap-frozen in liquid N_2_.

### Protein crystallization and data collection

Crystals of apo AtCKA3 developed in sitting drops composed of 0.4 μL of protein solution (AtCKA3 at 11 mg/mL in 20 mM HEPES [pH 7.5], 150 mM NaCl) and 0.4 μL of crystallization buffer (20 % [w/v] PEG 3,350, 0.2 M sodium malonate, 0.1 M Bis-Tris [pH 6.5]) suspended over

80 μL of the latter as reservoir solution. Crystals were transferred to reservoir solution supplemented with 20% (v/v) glycerol and snap-frozen in liquid N_2_. To obtain the AtCKA1 – InsP_6_ complex, AtCKA1 at 1 mg/mL in 20 mM HEPES (pH 7.5), 150 mM NaCl was mixed with InsP _6_ (Sigma P8810) to a final concentration of 5 mM and then concentrated to 20 mg/mL using an Amicon centrifugal filter with 10 kDa MWCO (Merck). Crystals grew in sitting drops (0.3 + 0.3 μL) in 10% (w/v) PEG 6,000, 0.1 M HEPES pH 6.5, and were cryoprotected by serial transfer in crystallization buffer supplemented with 20% (v/v) PEG 200. Crystals of the purified AtCKA1 – AtCKB1 complex in the presence of 5 mM InsP_6_ developed in hanging drops composed of 1 μL of protein solution (6 mg/mL in 20 mM HEPES [pH 7.5], 150 mM NaCl) and 1.0 μL of crystallization buffer containing 0.1 M Carboxylic acids (0.2 M sodium formate, 0.2 M ammonium acetate, 0.2 M sodium citrate tribasic dihydrate, 0.2 potassium sodium tartrate tetrahydrate, 0.2 M sodium oxamate), 0.1 M buffer system 2 (HEPES, MOPS, pH 7.2), 31% PrecipMix 2 (40% [v/v] Ethylene glycol, 20% [w/v] PEG 4,000, 20 % [w/v] PEG 3,350, 0.2 M sodium malonate, 0.1 M Bis-Tris [pH 6.5] (Morpheus G2, Molecular Dimensions) ^144^ suspended over 1 mL of the latter as reservoir solution. Crystals were frozen directly from the drop. All diffraction data were collected at beam line X06DA of the Swiss Light Source, Villigen, Switzerland. Data processing and scaling was done with XDS ^145^.

### Structure solution and refinement

The structures of apo AtCKA3 and the AtCKA1-InsP_6_ complex were solved by molecular replacement in Phaser^146^, using the previously reported AtCKA1 apo structure (PDB-ID: pdb_00006xx6) as search model^123^. The structures were completed in iterative rounds of manual model-building in COOT^147^ and restrained refinement in phenix.refine^148^. The refined structure of AtCKA1 and an AlphaFold model for AtCKB1 (AF-A0A5S9YCV4-F1) were used as search models to determine the structure of a the AtCKA1-AtCKB1 complex. The molecular replacement solution comprised a α_2_β_2_ hetero-tetramer, and the complex structure was completed by restrained NCS (non-crystallographic symmetry) refinement and using target restraints to the AtCKA1 structure and the ATCKB1 model in autoBUSTER (version 2.10.4)^149^. Inspection of the final models with phenix.molprobity^150^ revealed excellent stereochemistry (Supplementary Table 3). Structural representations were done with Pymol (https://sourceforge.net/projects/pymol/) and ChimeraX^140^.

### Grating-coupled interferometry

GCI experiments were performed on a CreoptixWAVE system (Malvern). After PCP chip conditioning, NeutrAvidin (Thermo Fisher) was immobilized by amine-coupling, followed by coupling of biotinylated 1/3-biotin-InsP_6_ (InsP_6_), 1/3-biotin-5PCP-InsP_5_ (InsP_7_) or 3-biotin1,5(PCP)_2_-InsP_4_ (InsP_8_) in 0.2x buffer with target level of 40 pp/mm^2^. Purified wild type and mutant AtCKA1 was used as analyte in 20 mM HEPES pH 7.5, 300 mM NaCl, 0.05% (w/v) BSA, 0.01 (v/v)% Tween-20 at a maximum concentration of 5 μM in five serial 1:2 dilutions at a flow rate of 100 μL/min at 25°C. All experiments were done in duplicates, blank injections were used for double referencing. Data analysis was performed with the Creoptix WAVEcontrol software (version: 4.7.2) and data were fitted with a bivalent model (two InsP_6_/PP-InsP binding sites). Data were plotted in R (https://www.r-project.org, version: 4.1.2).

### Radioactive kinase assay

All reactions were performed in 20 μL volume at room temperature (RT) for 1 h. 1 µM of purified wild-type or mutant AtCKA1 was incubated with 10 µM AtSOG1 or AtCCA1 substrate, or with 0.1 mg/mL dephosphorylated bovine milk α-casein (Sigma) in 50 mM HEPES pH 7.5, 300 mM NaCl, 1 mM MgCl2, 1 mM DTT. InsP_6_ was used at a final concentration of 200 µM, TTP-22 (Merck) at a final concentration of 5 µM. Reactions were initiated by the addition of 5 µCi radioactive ATP-γ ^32^P (Revvity). To maintain a stable ATP concentration, reactions were substituted with cold ATP (Sigma) to a final concentration of 50 μM. After 1h, reactions were stopped by adding 6xLämmli sample buffer and samples were boiled at 95°C for 5 min. The entire reaction was loaded onto 4-15% Mini-PROTEAN TGX gels (Bio-Rad) or 16.5% Mini-PROTEAN Tris-Tricine gels (Bio-Rad) and proteins were separated at a constant voltage of 80V in Tris-glycine (25 mM Tris, 192 mM glycine, 0.1% SDS, pH 8.3) running buffer for 50-60 min. The gels were exposed to a phosphoscreen (Fuji) for 1h, and the signal was read using the Typhoon biomolecular imager (Cytiva).

### Generation of transgenic Arabidopsis lines

The coding sequences of wild-type AtCKA3 wild-type, a kinase-dead version (K63M/D170N) and the 6xmut InsP_6_/PP-InsP binding site mutant (K69A,K71A,K72A,K242A,H271A,K274A) were synthesized (Twist bioscience) and cloned by Golden-Gate cloning into a vector providing a ubiquitin10 (proAtUBI10) promoter, a C-terminal 3xFlag (DYKDHD-G-DYKDHD-I-DYKDDDDK) tag, a NOS terminator and a FastRed selection marker. Constructs were introduced into *Agrobacterium tumefaciens* strain pGV3101 into the previously reported *A. thaliana cka1 cka2 cka3* (*cka1/2/3*) T-DNA insertion triple mutant^77^ using the floral dip method^151^. Transformants were identified by mCherry fluorescence with a Zeiss Axio Zoom.V16 stereo microscope (mRFP filter) and a HXP200C illuminator. Four stable homozygous, single insertion T3 lines were isolated for each construct.

The *cka4* and *ckb1/2/3/4* mutants were generated by clustered regularly interspaced palindromic repeats (CRISPR/Cas9) gene editing^152^. Guide-RNAs (gRNAs) were designed using the CRISPR-P v2 website (http://crispr.hzau.edu.cn/CRISPR2/) (Supplementary Table 4). Constructs were generated by annealing and ligating forward and reverse primers containing guides sequences to entry vectors resulting in U6::gRNA. These entry vectors were then used in Golden-Gate reactions to create final vectors containing U6::gRNA, proAtRPS5::Cas9 and FastRed selection. Constructs were introduced into Agrobacterium tumefaciens strain GV3101 and subsequently transformed into *A. thaliana* Col-0 ecotype using the floral dip method^151^. Transformants were selected by mCherry fluorescence. Genotyping of CRIPR-Cas9 events was done by PCR followed by Sanger sequencing, and Cas9 was segregated out in T2 generation. Homozygous quadruple *ckb1/2/3/4* and single *cka4* mutants homozygous were isolated in T4 generation by PCR and verified by Sanger-sequencing (Microsynth, AG) (Supplementary Fig. 7).

### Seedling root and hypocotyl length quantification

Seeds were gas sterilized, put on ^1/2^MS (Duchefa) plates and stratified for 48 h. Plates were then transferred to a Percival growth chamber with 16h / 8h light / dark cycle at 22°C for 10 d. 30 seedling were scanned on a flat bed scanner (Epson Perfection V600 Photo) and primary root length were manually measured in Fiji (version 1.54)^153^. For hypocotyl phenotyping seeds were gas sterilized, put on ^1/2^MS plates and stratified for 48 h. Plates were then wrapped in aluminum foil, moved to a Percival at 22°C and kept in the dark for 3 d. 30 seedling per line were scanned and hypocotyl length were manually measured using Fiji. In each case, a Dunnett test^154^ was performed to assess the statistical difference of the different genotypes compared to the Col-0 control (\**p* < 0.05, \*\*\**p* < 0.01, \*\*\**p* < 0.001).

### Flowering time assay

Seeds were gas sterilized, put on ^1/2^MS plates and stratified for 48 h. Plates were then moved to a Percival growth chamber with 16h / 8h light / dark cycle at 22°C for 7 d, after which 30-40 seedling per line were transferred to soil and kept in controlled environment with 16 h / 8 h light/dark regime at 22°C. Plants were scored for flowering by visible emergence of the inflorescence stem, and the number of leaves was counted at the same time. *35S*::*FT*^155^ and *cry2-1*^156^ were included as controls. Dunnett-type, two-sided multiplicity-adjusted *p*-values for the comparison against the Col-0 wild-type control are reported^154^. The discrete primary endpoints (days to flower and number of leaves) were modeled separately as over-dispersed Poisson-distributed counts using a generalized linear model approach within the framework of general parametric models^157^ as implemented in the CRAN package multcomp and sandwich in R (version 4.2.2)^158^ (\**p* < 0.05; \*\*\**p* < 0.001).

### Western blotting

∼100mg of Arabidopsis tissue per sample were harvested and snap-frozen in liquid N _2_ in 2 mL Eppendorf tube with glass beads. Samples were ground in a tissue lyzer (MM400, Retsch) and 300 µL of extraction buffer (50 mM Tris [pH 7.5], 150 mM NaCl, 1% (v/v) protease inhibitor cocktail [Sigma]) was added to each tube. Samples were gently agitated at 4°C for 20 min followed by centrifugation at 20,000 × g for 15 min at 4°C. 150 µL of supernatant was transferred to a new tube and 30 µl of 6xSDS loading buffer (0.5 M Bis-Tris [pH 6.6], 50% [v/v] glycerol, 6% [w/v] SDS, 0.5M DTT with 0.05% [w/v] bromophenol blue) was added. Samples were boiled at 95°C for 5 min and 30 μL were loaded on a 10% SDS-PAGE gel. Proteins were transferred to a nitrocellulose membrane (0.2 μm, GE Healthcare) for 2 h at 0.3 A. Membranes were then blocked for 1 h with 5% (w/v) milk powder in TBS-T (50 mM Tris pH 7.5, 150 mM NaCl, 0.1% [v/v] Tween 20) at RT. Membranes were incubated at RT for 2 h with Anti-FLAG M2-Peroxidase (Sigma, 1:5,000 dilution). Membranes were washed three times with TBS-T and were detected using SuperSignal™ West Femto Maximum Sensitivity Substrate (Thermo Scientific) and visualized with Amersham Imager (Cytiva).

Marchantia proteins were extracted from 5-week-old thalli. About 200 mg of sample were collected in 2 ml Eppendorf tubes and homogenized in a tissue lyzer (MM400, Retsch) in liquid N_2_. Then, 300 µL of 1x PBS buffer supplemented with the plant-specific protease inhibitor cocktail (PE0230, Merck) was added and incubated for 30 min at 4 °C in an orbital shaker (Intelli-Mixer RM-2M, ELMI). The samples were then centrifuged at 10,000 × g for 20 min at 4 °C. Then, 150 µL of the supernatant was transferred in a new tube containing 30 µL of 6xSDS loading buffer and incubated at 95 °C for 5 min. A 10% BIS-Tris acrylamide SDS-PAGE was run with MES running buffer. Proteins were transferred to a 0.2 µm PVDF membrane (IB34002, Thermo Scientific) using a semi-dry protocol (iBlot3, Thermo Scientific). Membranes were blotted with TBS-Tween (0.1% [v/v]) – milk (5% [w/v) for 1 h at RT. For TagRFP detection, membranes were incubated with a rabbit polyclonal pan-Red Fluorescent Protein antibody (#pabr1, Chromotek) using a dilution of 1:5,000. Goat Anti-Rabbit IgG–HRP (A6154, Sigma) was used as secondary antibody in 1:10,000 dilution and incubated for 1 h at RT. Detection was performed using SuperSignal West Femto Maximum Sensitivity Substrate (34095, Thermo Scientific) and visualized with Amersham Imager (Cytiva).

### Yeast complementation assay

The *Saccharomyces cerevisiae* strains, plasmids and primers used in this study are listed in Supplementary Table 4. Preparation of media, yeast transformations, and genetic manipulations were performed according to established protocols. All recombinant DNA techniques were performed according to established procedures using Escherichia coli TOP10 cells for cloning and plasmid propagation. Mutations were generated with the QuickChange site-directed mutagenesis kit (Agilent Technologies, Santa Clara, CA, USA). All cloned DNA fragments and mutagenized plasmids were verified by Sanger-sequencing (Microsynth, AG). Centromeric plasmids encoding Cka1, Cka2, Cka1^KE^ (K75E, K77E, K78E) and Cka2^KE^ (K85E, K87E, K88E) were transformed into cka1Δcka2Δ pRS416-CKA2 shuffle strain^159^. The ability of Cka1^KE^ and Cka2^KE^ variants to complement the lethality of cka1Δcka2Δ strain was scored by spotting the transformants in 10-fold serial dilutions on synthetic deficient (SD) media plates lacking leucine (Leu) and containing 5-Fluoroorotic acid (5-FOA).

### SEC-MALS analysis

Experiments were carried out on an OmniSEC (Malvern). A Superdex 200 Increase 10/300 GL (Cytiva) was equilibrated in filtered and degassed buffer (20 mM HEPES pH 7.5, 300 mM NaCl, 0.02% (w/v) NaN_3_) and 1 mM InsP_6_, if applicable, on an Akta PURE system. After transfer to the OmniSEC RESOLVE, the system was further equilibrated in buffer until it a stable baseline was achieved. All runs were performed at a flow rate of 1 mL/min, at 25°C with injections of 50 µL of protein at a concentration of 5 mg/mL. A BSA calibration was performed before and after all measurements. Data were analyzed using the OMNISEC software as provided by the manufacturer (version: v10.41).

### Anthocyanin quantification in Arabidopsis

Seeds were gas sterilized, put on ^1/2^MS (Duchefa) plates and stratified for 48h. Plates were moved to a Percival at 22°C with 16h / 8h light / dark cycle for 10d. ∼20mg of seedling shoot material was placed into 5 ml of extraction buffer (18% [v/v] 1-propanol, 1% [v/v] HCl, 81% [v/v] H _2_O), boiled for 3 min at 95°C and incubated in the dark overnight at RT. Then the supernatant was cleared by centrifugation at 20,000 × g for 5 min at RT. The absorbance at 535 nm and 650 nm was measured using a spectrophotometer (Tecan Infinite M Nano). Anthocyanin levels were expressed as (A_535nm_–A_650nm_) g^−1^ fresh weight (FW). The Dunnett test was used to assess the statistical difference of the different genotypes compared to the Col-0 control (\*\*\**p* < 0.001)^154^.

### Quantification of seed abortion phenotype

*cka1 cka2 cka3 cka4*(-/+) plants were grown in soil for 5 weeks with a photoperiod of 16 / 8 hr at 22 °C. Seed abortion was evaluated on dissected siliques were aborted seeds are typically white and viable seeds green allowing the distinction between the two states. Pictures from 6 siliques from *cka1 cka2 cka3 cka4*(-/+) and Col0, respectively were taken with a binocular Nikon SMZ18 and analyzed for quantification in Fiji.

### Marchantia polymorpha growth conditions

Wild type and transgenic *M. polymorpha* plants were propagated through gemmae on ^½^Gamborg B5 medium (Sigma) adjusted to pH 5.5 with KOH, under constant LED-source white light (60 μmol/m^2^/s) at 22°C in 90 mm Petri dishes containing 1.2% (w/v) plant cell culture agar (Huber lab). Induction of the female and male sexual structures was done by transferring plants to soil conditions on pots covered by a transparent lid to maintain the humidity. Plants were maintained in the same light conditions supplemented additionally with far red light to stimulate the induction of the sexual structures. Fertilization was archived using water to release the sperms from the male and then inoculate them in the female structure using a micropipet. Spores were sterilized with 1 ml Milton solution for 20 min and spread in the regular ^½^B5 medium for one week in the same growing conditions. Sterilization solution was prepared by solubilizing one sterilizing tables (Milton 931011) in 500 mL of deionized H_2_O^160^.

### Generation of CKA and CKB knock-out and over-expression lines in Marchantia

Mpcka^ge^ and Mpckb^ge^ loss-of-function mutants were generated by CRISPR/Cas9 gene editing, as previously described^20^. gRNAs were designed using Casfinder (https://marchantia.info/tools/casfinder/). Two target sequences (Supplementary Table 4) were selected each and primers including forward 5’ CTCG-3’ and reverse 5’ AAAC-3’ overhangs, respectively were synthesized (Microsynth). Annealed primers were cloned into a pMpGE_En03 (Addgene #71535), plasmid digested with BsaI using T4 ligase (NEB). The gRNAs were subcloned into binary vector pMpGE010 (Addgene #71536) containing the Cas9 enzyme and a Hygromycin resistance gene as selection marker by Gatway Cloning, and transformed into *Agrobacterium tumefaciens* strain GV2260. Plant transformations were done using the spore transformation method^161^. Transgenic lines were genotyped by PCR using KOD polymerase (Merck) and Sanger sequencing, using primers flanking the target regions. We selected lines bearing insertions or deletions that lead to frame shifts and early stop codons (Supplementary Fig. 7). For generation of over-expression lines, MpCKA^wild^ ^type^, MpCKA^kinase-dead^ and MpCKA^6xmut^ were synthesized (Twist Biosciences) and sub-cloned in pDONR207 using Gateway cloning. Entry vectors were introduced into the binary plasmid pMpGWB-327 under the expression of mpEF1 promoter fussed to a C-terminal RFP and Chlorsulfuron as selection marker ^162^. Sex determination was done by PCR using specific primers to amplify fragments from the female and male sexual chromosome, respectively (Male_Fw CCAAGTGCGGGCAGAATCAAGT, Male_Rv TTCATCGCCCGCTATCACCTTC, Female_Fw GACGACGAAGATGTGGATGAC and Female_Rv GAAACTTGGCCGTGTGACTGA).

### Thallus area quantification

Thallus surface areas of different plant genotypes were quantified rom single plants grown from gemmae (1 gemmae per one round 90 mm petri dish) grown in ^½^B5 medium in time course experiments (n=12 per genotype). Top view images were acquired weekly using a flatbed scanner (Epson Perfection V600 Photo) and thallus areas were manually segmented in Fiji ^153^. Images were converted to 8 bit followed by manual selection of individual plants. Contrast was adjusted, and the thallus surface was selected using the threshold. The scale (2832 px to 9 cm) was set before quantification. Finally, the area of the segmented image was determined using the Particle Quantify function as implemented in Fiji. A Dunnett test was used to assess the statistical difference of the different genotypes compared to the Tak-1 wild type (\*\*\**p* < 0.001) or to the *proMpEF1*::*MpCKA*^wild^ ^type^-RFP control (^###^*p* < 0.001).

### UV-B-absorbing compounds extraction and quantification in Marchantia

For each sample 50 mg of tissue from 5 weeks old plants was collected in an Eppendorf tube and frozen in liquid N_2_. The tissue was homogenized using a tissue lyzer (MM400, Retsch) at 30hz for 15 sec. Then, 500 µL of extraction buffer (80% [v/v] methanol and 1% [v/v] HCl) was added. Samples were mixed and incubated overnight at 4°C. Samples were spun down at 20,000 × g for 3 min at RT. Absorbance at 310 nm was measured using a spectrophotometer (Tecan Infinite M Nano) and normalized to the sample fresh weight (FW)^127^. A Dunnett test was used to assess the statistical difference of the different genotypes compared to the Tak-1 wild type (\*\*\**p* < 0.001), or to *proMpEF1*::*MpCKA^wild^ ^type^-RFP* (^###^*p* < 0.001)^154^.

## Supporting information

alphafold and boltz-2 models

## Figure legends

**Fig. S1:**
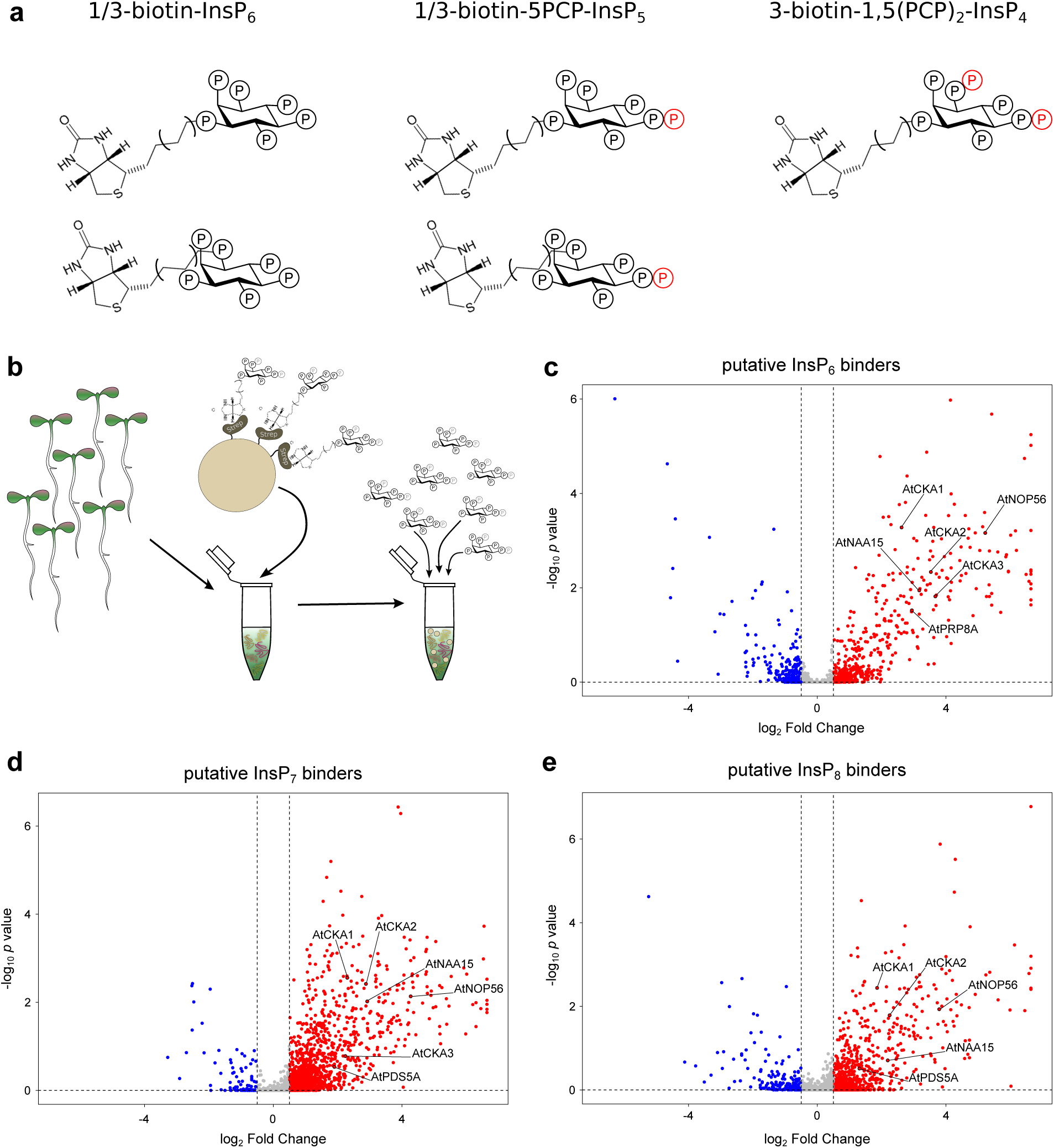
A proteomic screen identifies novel InsP_6_/PP-InsP binding proteins. **a** Chemical structures of the biotinylated InsP_6_, InsP_7_ and InsP_8_ analogs used for the screen. **b** Schematic overview of the PP-InsP pull down experiment using streptavidin-coupled sepharose bead-adsorbed InsP_6_/PP-InsP analogs. Bound target proteins were eluted by competition with soluble InsP_6_, or InsP_7_/InsP_8_ analogs, respectively. Volcano plots showing significantly enriched proteins (FDR < 0.05) in the **c** InsP_6_, **d** InsP_7_ and **e** InsP_8_ elutions. Selected proteins discussed in the text are indicated.

**Fig. S2:**
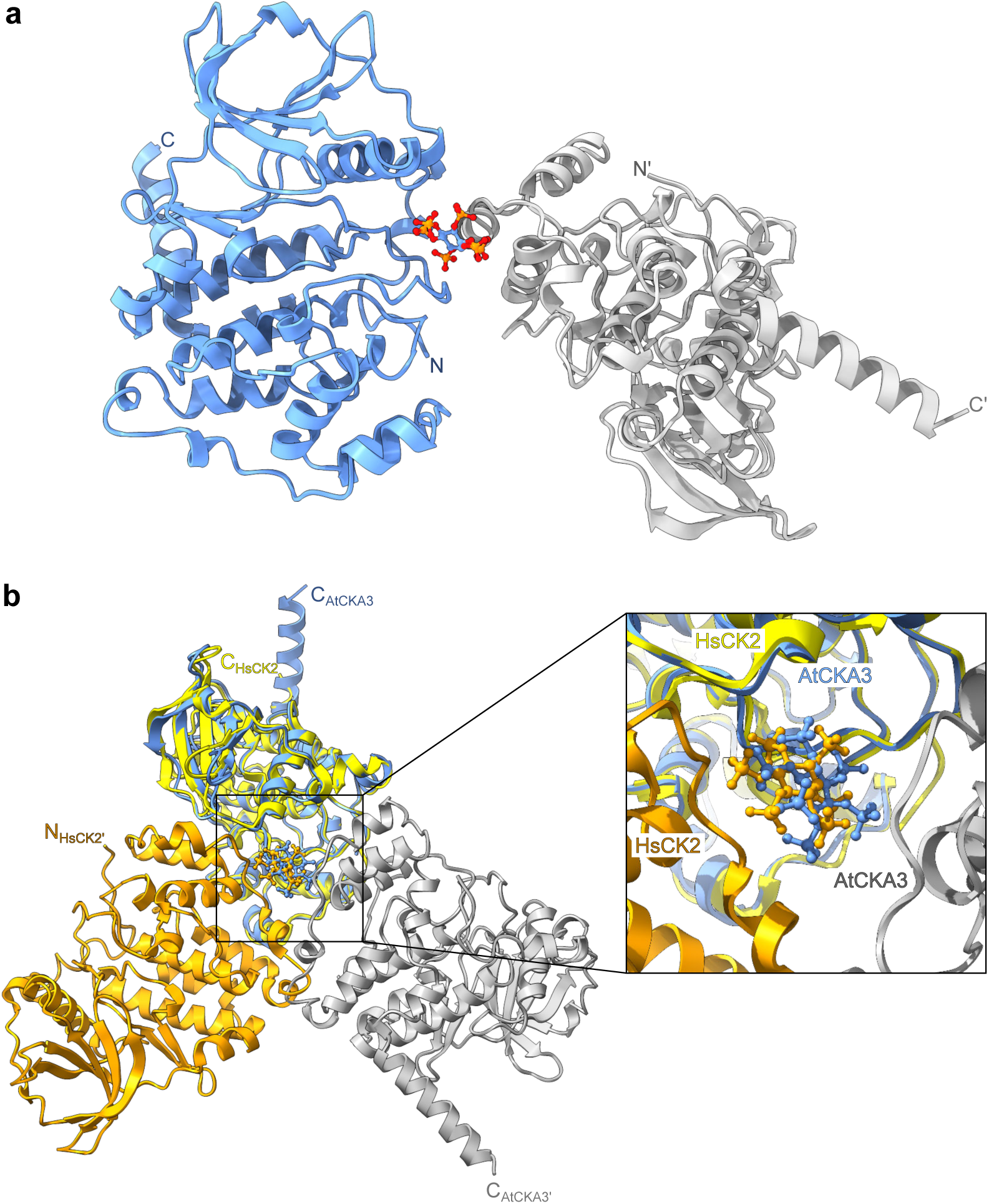
The InsP_6_ ligand is sandwiched between two symmetry-related CK2 molecules in the AtCKA1 and HsCK2 complex structures. **a** Ribbon diagram of AtCK2 (in blue) in contact with a symmetry-related molecule (in gray). The InsP_6_ (in bonds representation) at the interface between the two molecules is coordinated by residues from the basic binding surface near the nucleotide-binding cleft, as well as from the C-lobe of the kinase. **b** Structural superposition of the AtCKA1 – InsP_6_ complex (colors as in panel A) with the previously reported HsCK2 – InsP_6_ complex (in yellow and orange, respectively, PDB-ID pdb_00003w8l, r.m.s.d is ∼ 1 Å comparing 323 corresponding C_α_ atoms) reveals highly similar InsP_6_ (in bonds representation) interactions.

**Fig. S3:**
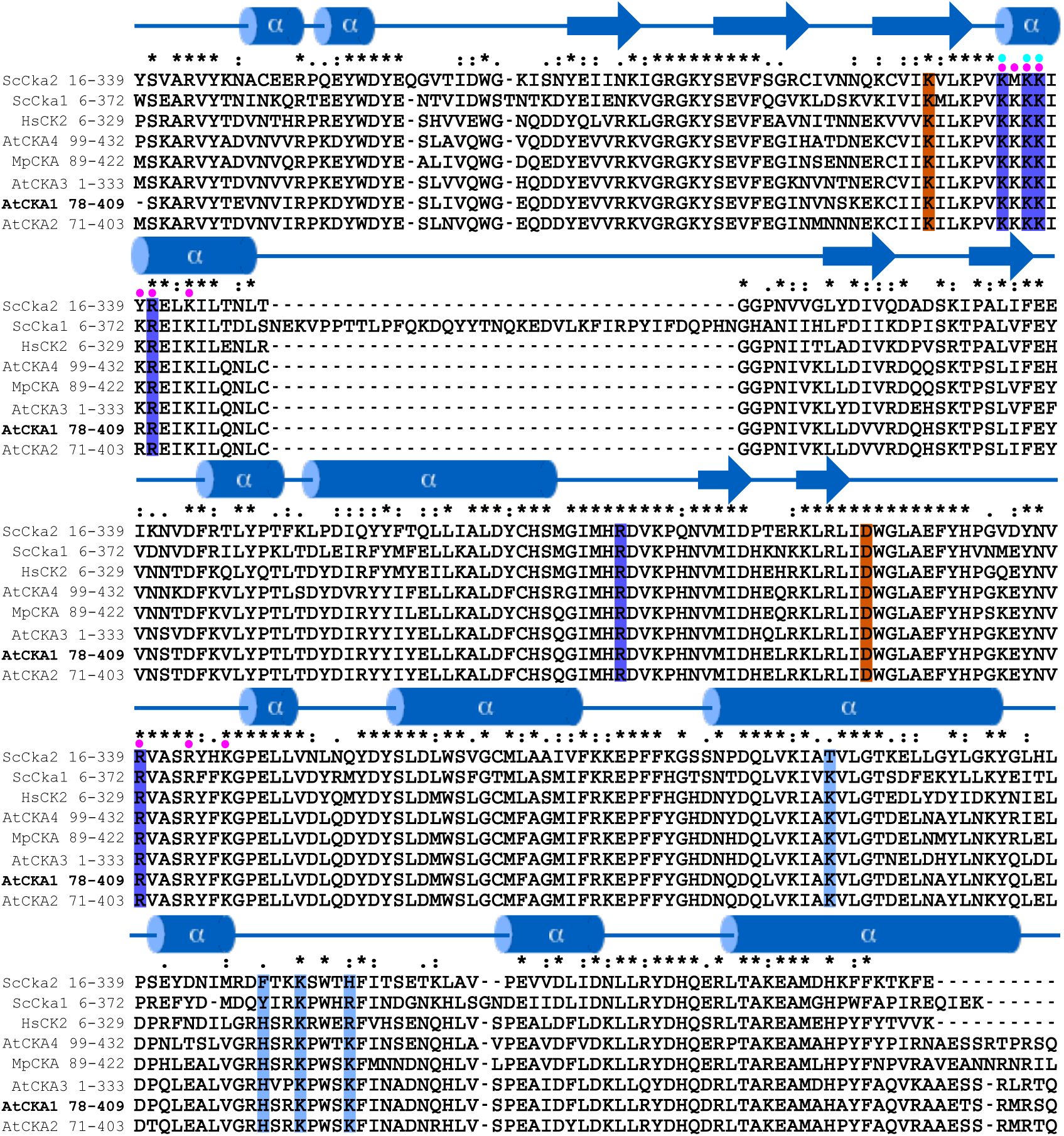
The InsP_6_/PP-InsP binding surfaces are conserved among CK2 α-subunits from different organisms. Multiple sequence alignment of protein kinase catalytic subunits ScCKA2 (UniProt ID: P19454), ScCKA1 (P15790), HsCK2 (P68400), AtCKA4 (O64816), MpCKA (A0A2R6WKI0), AtCKA3 (O64817), AtCKA1 (Q08467), and AtCKA2 (Q08466), generated using MUSCLE^163^. The alignment includes the secondary structure assignment of AtCKA1 as determined by DSSP^164^. Residues mutated to generate kinase-dead variants of AtCKA1 and AtCKA3 are highlighted in orange. Residues contributing to the InsP₆/PP-InsP binding surface located near the nucleotide-binding cleft or within the C-terminal lobe are shown in dark blue and light blue, respectively. Conserved basic residues previously mutated in HsCK2 are indicated by a magenta dot^62^. The InsP_6_/PP-InsP binding site mutations in ScCka1^KE^ (K75E,K77E,K78E) and ScCka2^KE^ (K85E,K87E,K88E) are highlighted by a cyan dot.

**Fig. S4:**
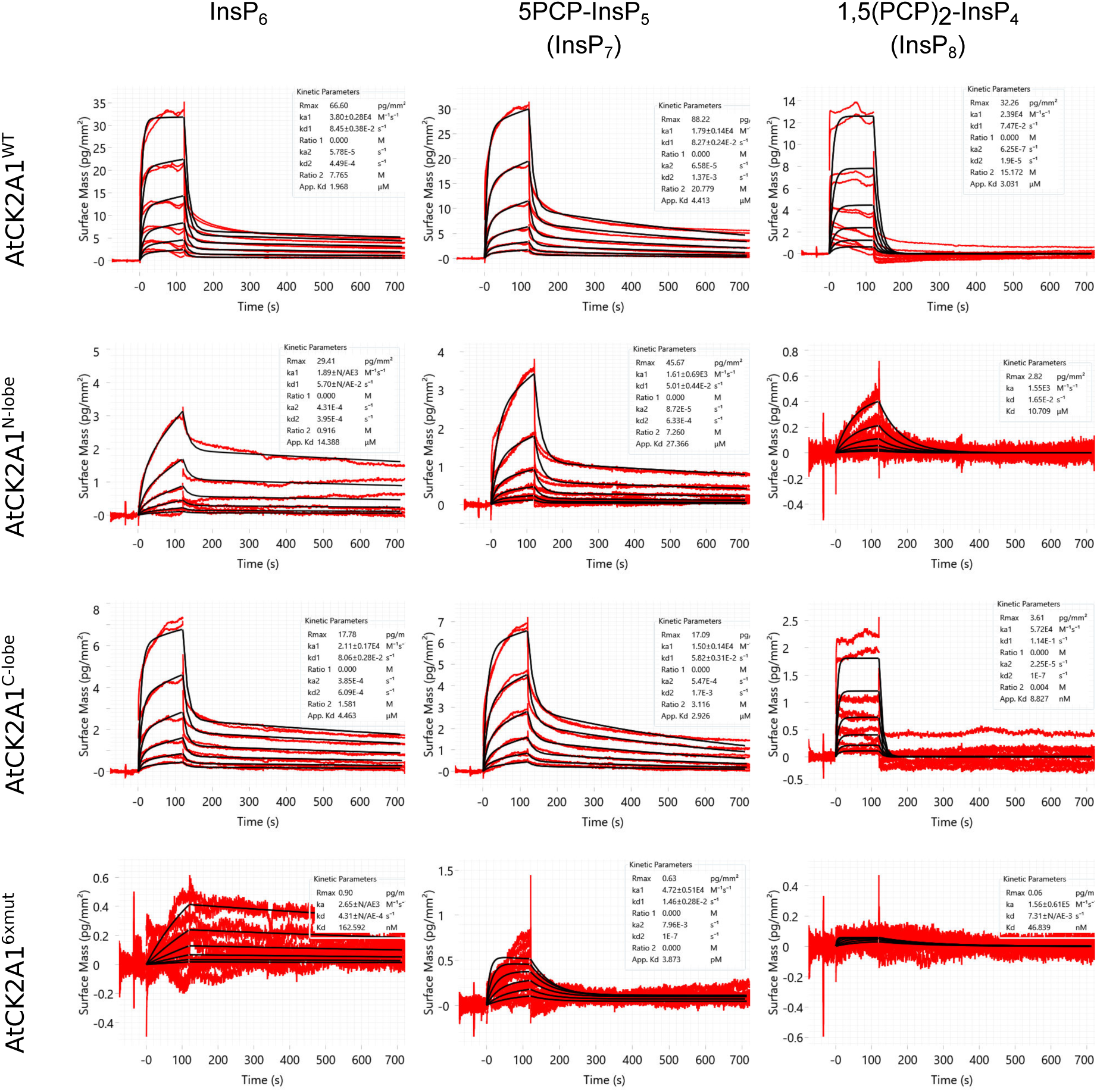
Grating coupled interferometry sensorgrams related to Fig 4f. GCI analyses of ligand-binding kinetics between wild-type or mutant AtCKA1 vs. 1/3-biotin-InsP_6_ (InsP_6_), 1/3-biotin-5PCP-InsP_5_ (InsP_7_) or 3-biotin1,5(PCP)_2_-InsP_4_ (InsP_8_) coupled to the GCI chip. Shown are sensorgrams (n=2, red); global fits are in black.

**Fig. S5:**
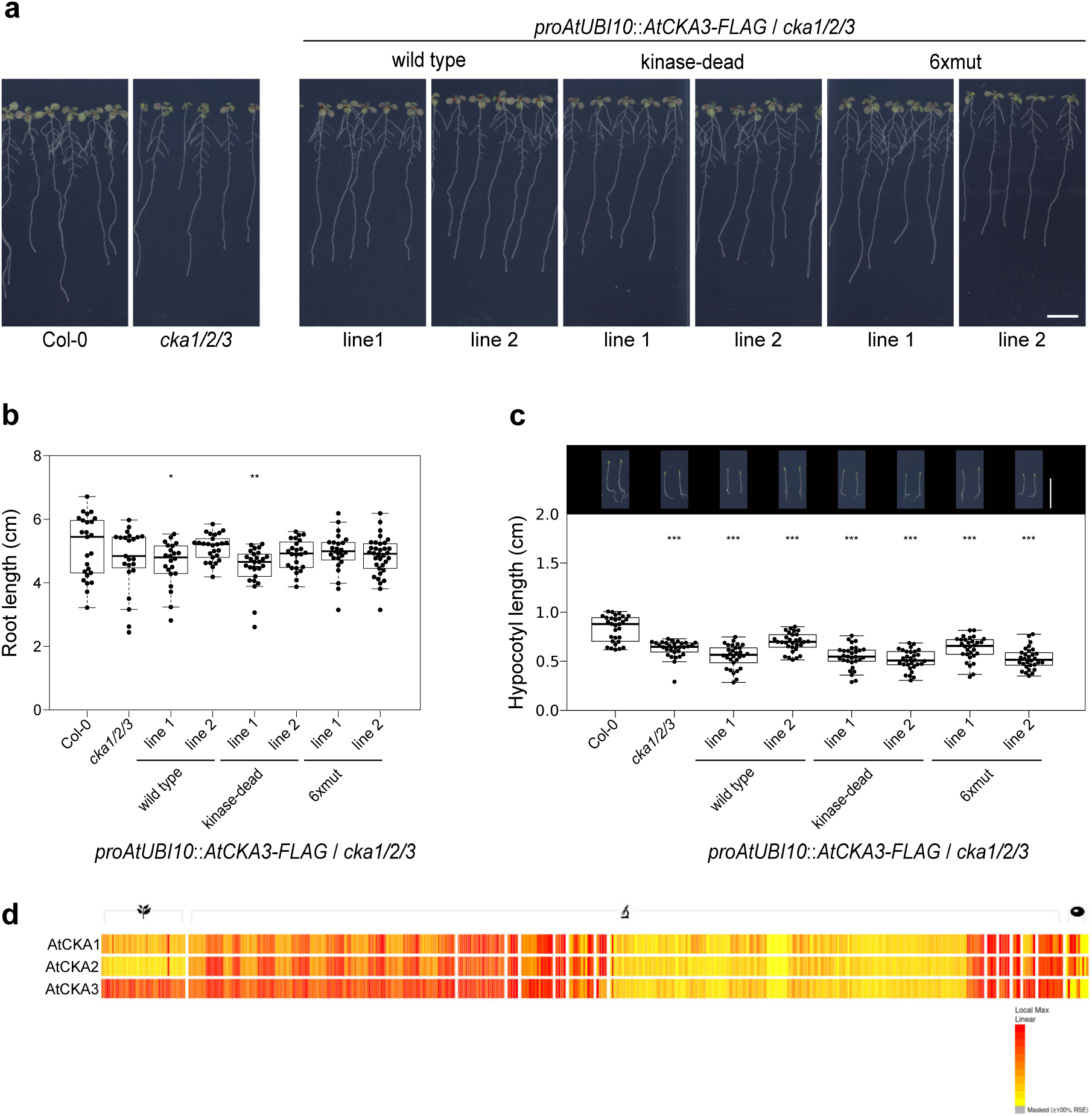
The *cka1/2/3* root and hypocotyl phenotypes cannot be robustly rescued by constitutive expression of AtCKA3. **a** Root phenotypes of 14 d-old seedlings grown on ^1/2^MS plates at 22°C (n=30; scale bar = 1cm). **b** Box plots representing primary seedling root length of the lines shown in **a**. Bold black line, median; box, interquartile range (IQR); whiskers, lowest/highest data point within 1.5 IQR of the lower/upper quartile. Dunnett-type, two-sided multiplicity-adjusted *p*-values for the comparison against the Col-0 control are shown alongside (\**p* < 0.05; \*\**p* < 0.01; \*\*\**p* < 0.001). **c** Hypocotyl phenotypes (top) and hypocotyl length quantification (bottom) of 5 d-old dark-grown seedlings (scale bar = 1cm). **d** Comparative expression heat map for AtCKA1 (At5g67380), AtCKA2 (At3g5000) and AtCKA3 (At2g23080) calculated with the BAR ePlant Heat Map viewer (https://bar.utoronto.ca/eplant/)^165^.

**Fig. S6:**
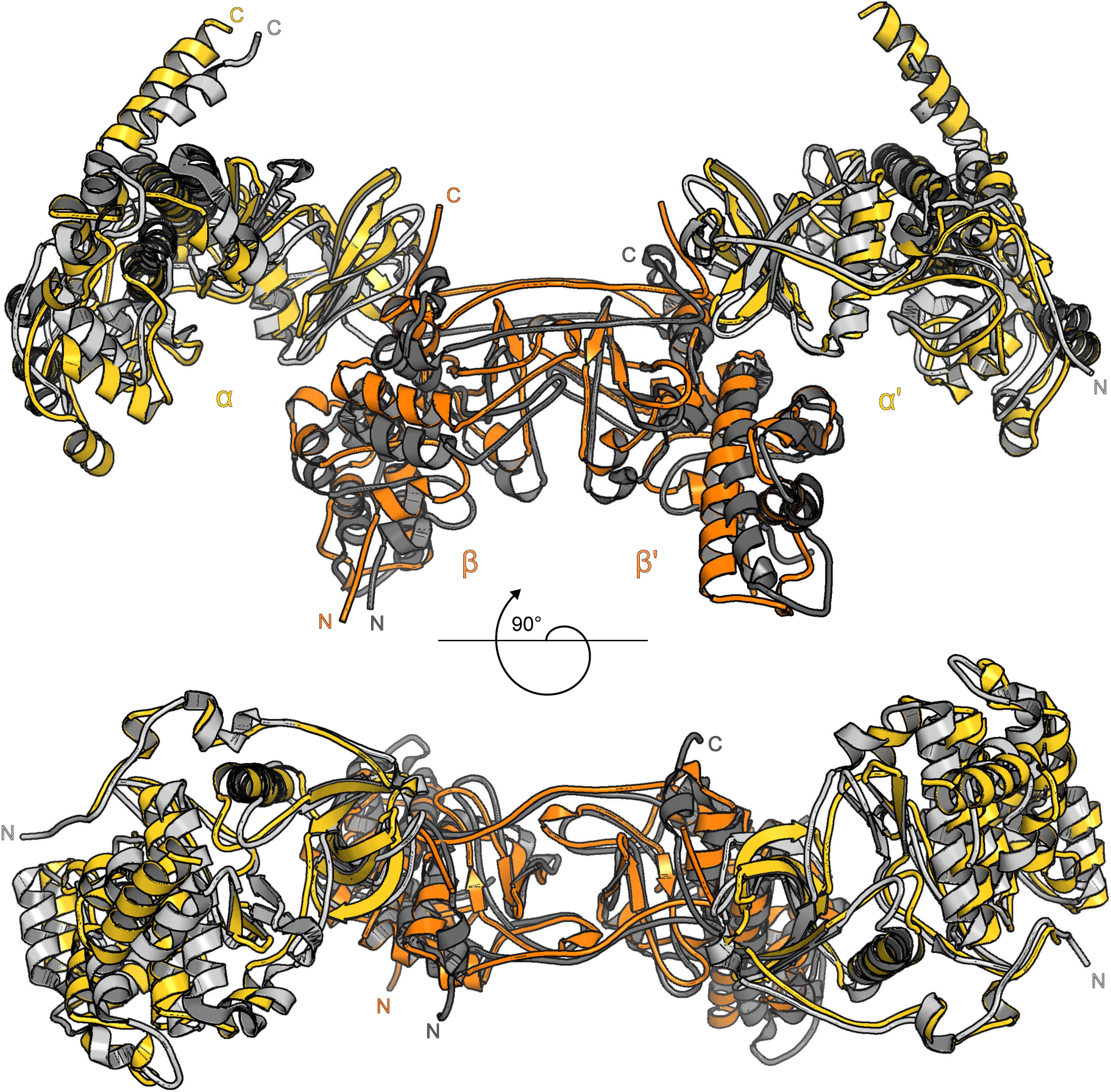
The Arabidopsis and human CK2 holoenzymes are structurally related. Structural superposition (r.m.s.d. is ∼ 3.9 Å comparing 891 corresponding C_α_ atoms) of the AtCKA1 – AtCKB1 complex (shown in yellow and orange, respectively) and the previously reported human holoenzyme (PDB-ID: pdb_00001jwh; in gray)^59^.

**Fig. S7:**
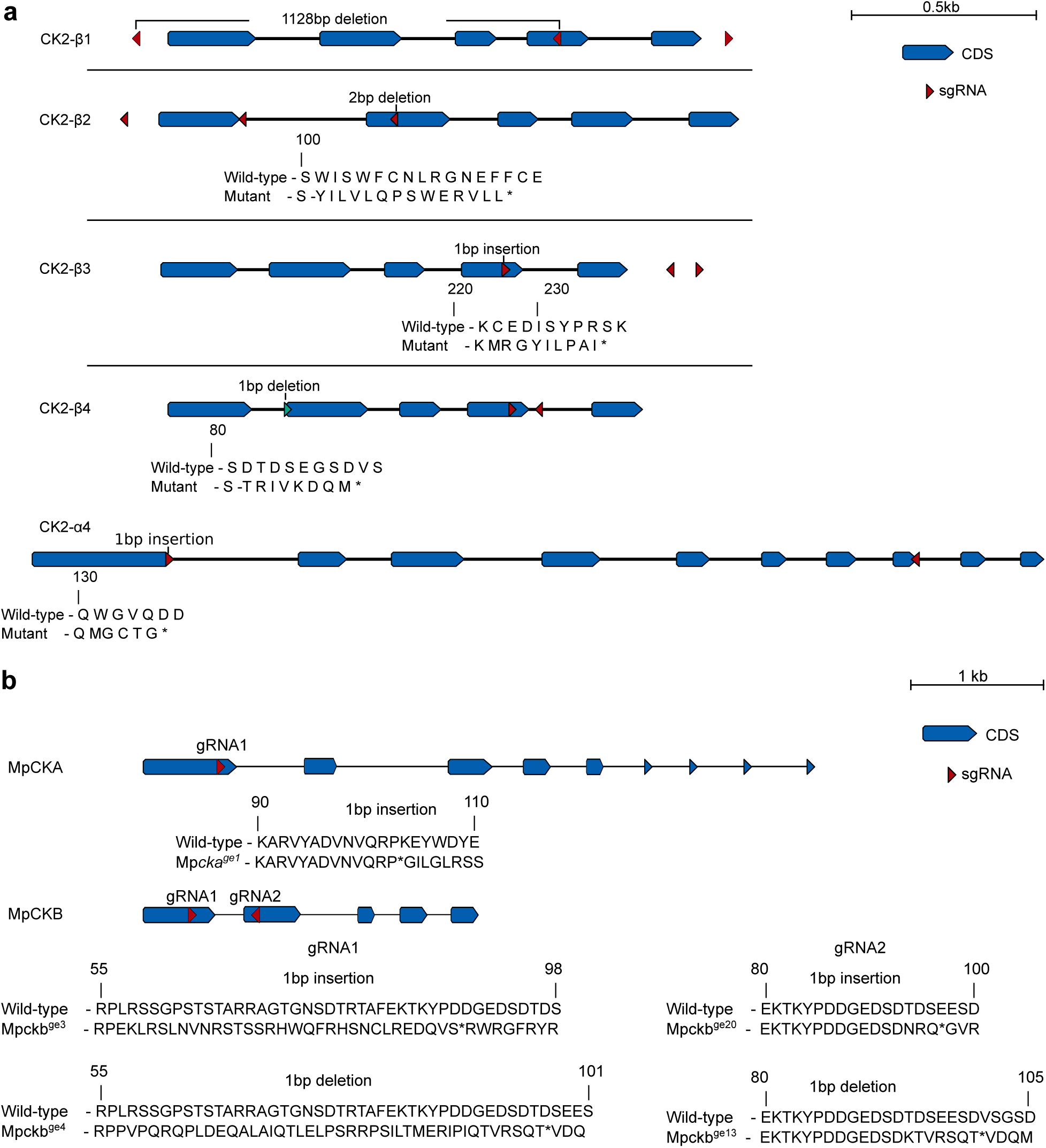
CRISPR/Cas9 gene editing events for the Arabidopsis and Marchantia mutants generated for this study. Schematic overview of the **a** AtCKB1, AtCKB2, AtCKB3, AtCKB4, AtCKA4 and **b** MpCKA and MpCKB genes with exons depicted in blue and introns as lines. CRISPR-Cas9 sgRNA guide sequences are represented as red arrowheads. All CRISPR events are shown below the gene targeted region, as confirmed by Sanger sequencing.

## Acknowledgments

We thank Chentao Lin for the *cka1/2/3* mutant, Markus Schmid for providing us with the *35S*::*FT* seeds, Roman Ulm for the *cry2-1* allele, Zhoubo Hu for the *35S*::*YFP* control line, the Proteomics Core Facility, Centre Medical Universitaire (CMU), University of Geneva for mass spectrometry analysis, and the staff of the Swiss Light Source (SLS) for technical assistance during data collection. We thank D. Couto for his input during the initial stage of this project, A. Caregnato for help with plotting data and R. Ulm, J. Santiago, E. Levy and P. Rieu for critically reading the manuscript. This work was supported by the HORIZON EUROPE European Research Council consolidator grant 818696 INSPIRE (to M.H.), by Swiss National Science Foundation Sinergia grant CRSII5_209412 (to M.H. D.F. and V.G.P.), and by a International Research Scholar Award 55008733 by the Howard Hughes Medical Institute (to M.H.).

## Disclosures

The authors declare no competing interests.

## Data availability

The mass spectrometry proteomics data have been deposited to the PRIDE/ProteomeXchange database (https://www.ebi.ac.uk/pride) with accession numbers PXD063319 (InsP_6_/InsP_7_/InsP_8_) and PXD069954 (AtVIH2-mCit). Crystallographic coordinates and associated structure factors have been deposited with the Protein Data Bank (http://rcsb.org) with accession numbers pdb_00009TFW (AtCKA3 apo), pdb_00009TFX (AtCKA1 – InsP_6_) and pdb_00009TFY (AtCKA1 – AtCKB1 complex).

## Author contributions

K.S. performed the PP-InsP interaction screen with material prepared by A.R.. K.S. cloned, expressed and purified proteins, performed binding and kinase assays, SEC-MALS experiments and crystallized proteins. K.S. and M.H. solved, build and refined the structures, O.P-T. with the help of

L.B. and F.R-R. generated transgenic Arabidopsis lines, performed flowering time assays, hypocotyl and root growth assays and western blotting, F.R-R. generated transgenic Marchantia lines, performed phenotyping experiments in Marchantia and in Arabidopsis and together with H.C. performed segregation analyses. Y.V. performed the yeast complementation assays. H.C. performed the AtVIH2 IP/MS experiment and split-luciferase assays. L.A.H. analyzed data and performed statistical analyses. D.F., V.G.P. and M.H. acquired funding and analyzed data. M.H. conceived the project and wrote the manuscript, with input from all authors.

---

**Table S3:**
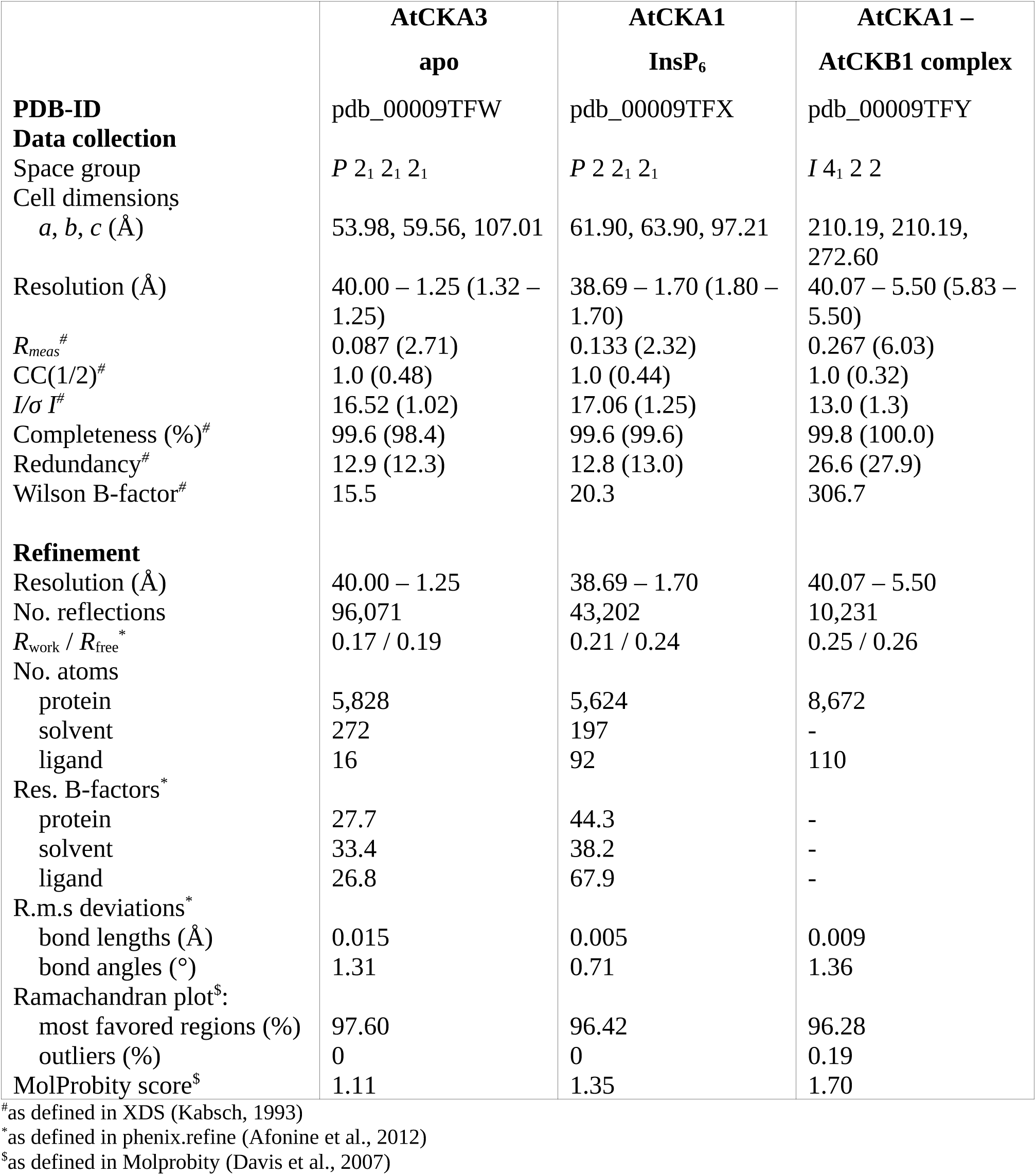
Crystallographic data collection and refinement statistics.

## Notes

### Competing Interest Statement

The authors have declared no competing interest.

